# Disease modeling with human neurons reveals LMNB1 dysregulation underlying DYT1 dystonia

**DOI:** 10.1101/2020.08.11.246371

**Authors:** Baojin Ding, Yu Tang, Shuaipeng Ma, Masuma Akter, Meng-Lu Liu, Tong Zang, Chun-Li Zhang

## Abstract

DYT1 dystonia is a hereditary neurological disease caused by a heterozygous mutation in *torsin A* (*TOR1A*). While animal models provide insights into disease mechanisms, significant species-dependent differences exist since mice with the identical heterozygous mutation fail to show pathology. Here, we model DYT1 by using human patient-derived motor neurons. These neurons with the heterozygous *TOR1A* mutation show markedly thickened nuclear lamina, disrupted nuclear morphology, and impaired nucleocytoplasmic transport, whereas they lack the perinuclear “blebs” that are often observed in animal models. Importantly, we further uncover that the nuclear lamina protein LMNB1 is specifically dysregulated in expression and subcellular localization. LMNB1 downregulation can largely ameliorate all the cellular defects in DYT1 motor neurons. These results reveal the value of disease modeling with human neurons and provide novel molecular mechanisms underlying DYT1 dystonia and potentially other neurological diseases with impaired nucleocytoplasmic transport.

## Introduction

Dystonia is a common movement disorder characterized by sustained or intermittent muscle contractions causing abnormal movements and/or postures ^1^. The clinical characteristics and underlying causes of dystonia are very heterogeneous, so the cellular mechanism for dystonia remains elusive ^2^. Childhood-onset torsin dystonia, also called DYT1 dystonia, represents the most frequent and severe form of hereditary isolated dystonia ^1,3^. Most cases of typical DYT1 dystonia are caused by a heterozygous 3-bp deletion (ΔGAG) in exon 5 of the *TOR1A* gene, resulting in the loss of one of the two adjacent glutamate residues (E302 or E303) in the carboxy-terminal region of torsin A protein (TOR1AΔE) ^4,5^.

Protein TOR1A belongs to the superfamily of AAA+ (ATPases associated with diverse cellular activities) proteins ^4^. It is widely distributed throughout the human central nervous system (CNS) ^6^. Most TOR1A proteins are embedded in the lumen of the endoplasmic reticulum and the endomembrane space of the nuclear envelope (NE) ^7^. Accumulating evidence indicates that TOR1A plays an important role in membrane-cytoskeleton interactions ^8^. At the NE, TOR1A is associated with the linkage between NE and cytoskeleton *via* binding the KASH domain of nesprins ^9^. It is also involved in the regulation of NE membrane morphology ^10–12^, the localization and function of nuclear pore subunits ^13^, and the nuclear egress of large ribonucleoprotein granules ^14^. Furthermore, TOR1A regulates neurite extension ^15^ and synaptic vesicle recycling ^16^, underscoring its critical functions in neuronal development and function.

Although TOR1AΔE has been shown to be a loss-of-function mutation ^10,17,18^, the cellular mechanisms underlying DYT1 dystonia remain largely unknown. Traditionally, dystonia is considered as a disorder of the basal ganglia ^19^, supported by the findings in *Tor1a* mutant mice including the early structural and functional plasticity alterations in the corticostriatal synapses ^20^ and the dysregulation of the striatal dopaminergic neutrons ^21^. The cerebellum *via* direct modulation of the basal ganglia is also shown as a major site of dysfunction in *Tor1a* knockout mice ^19,22^. One cellular hallmark in *Tor1a* knockout mice is the abnormal NE morphology in multiple areas of the CNS including the large ventral horn of the spinal cord ^10^, indicating that the lower motor neurons might be particularly affected. Unlike the majority of human DYT1 patients who are affected by the heterozygous *TOR1A* mutation, mice with the identical mutation as a heterozygote nonetheless failed to exhibit any pathological phenotypes ^10^, clearly indicating there might be species-dependent differences on disease mechanisms. However, DYT1 has never been previously modeled in human neurons.

Through cell fate reprogramming, we successfully generated human motor neurons (MNs) from fibroblasts of DYT1 patients and healthy controls *via* two strategies: direct conversion (diMNs) and differentiation through induced pluripotent stem cells (iPSCs) (iPSC-MNs). We found that DYT1 neurons generated by both methods show impairments in neuron development, NE morphology, mRNA nuclear export, and protein transport. We further discovered that nuclear lamin B1 (LMNB1) is dysregulated in both expression and subcellular localization in patient MNs. Importantly, LMNB1 downregulation can largely ameliorate defects of NE morphology and nuclear transport in patient neurons. These findings indicate that LMNB1 dysregulation may contribute to DYT1 dystonia and provide a novel molecular target for intervention.

## Results

### Directly Induced MNs from Fibroblasts of DYT1 Patients

Because dystonia is a movement disorder and the ventral horn neurons in the spinal cord are most severely affected in *Tor1a* mutant mice ^10^, we first generated MNs from human fibroblasts using a direct reprogramming method as we have previously described ^23,24^ (Fig. 1A). The fibroblasts of four DYT1 patients and four age-matched healthy controls were used for this study (Table S1). Although sex does not affect the reprogramming efficiency ^25^, all healthy controls were selected to be sex-matched to patient cell lines. Transduction with reprogramming factor-expressing lentivirus enabled adult human fibroblasts to gradually become neuron-like and grow complex neurites with time (Fig. S1). Immunocytochemistry confirmed their expression of the neuronal marker TUBB3 (Fig. 1B) and early MN marker HB9 (Fig. 1C) ^26^. Neuronal conversion efficiency was similar among all these fibroblasts, with about 85% of virus-transduced GFP+ cells as TUBB3+ neurons (Fig. 1D). Approximately 80% of these neurons showed strong HB9 expression and there was no difference between different cell lines (Fig. 1E). At later stages, such as 7 or 9 weeks post viral infection (wpi), reprogrammed neurons expressed high levels of choline acetyltransferase (CHAT) and exhibited discrete puncta of the pre-synaptic marker synapsin 1 (SYN1) (Fig. 1F, G). Thus, fibroblasts from DYT1 patients as well as healthy controls could be efficiently and directly converted into mature MNs, which were hereafter referred as diMNs (directly induced motor neurons).

**Figure 1.**
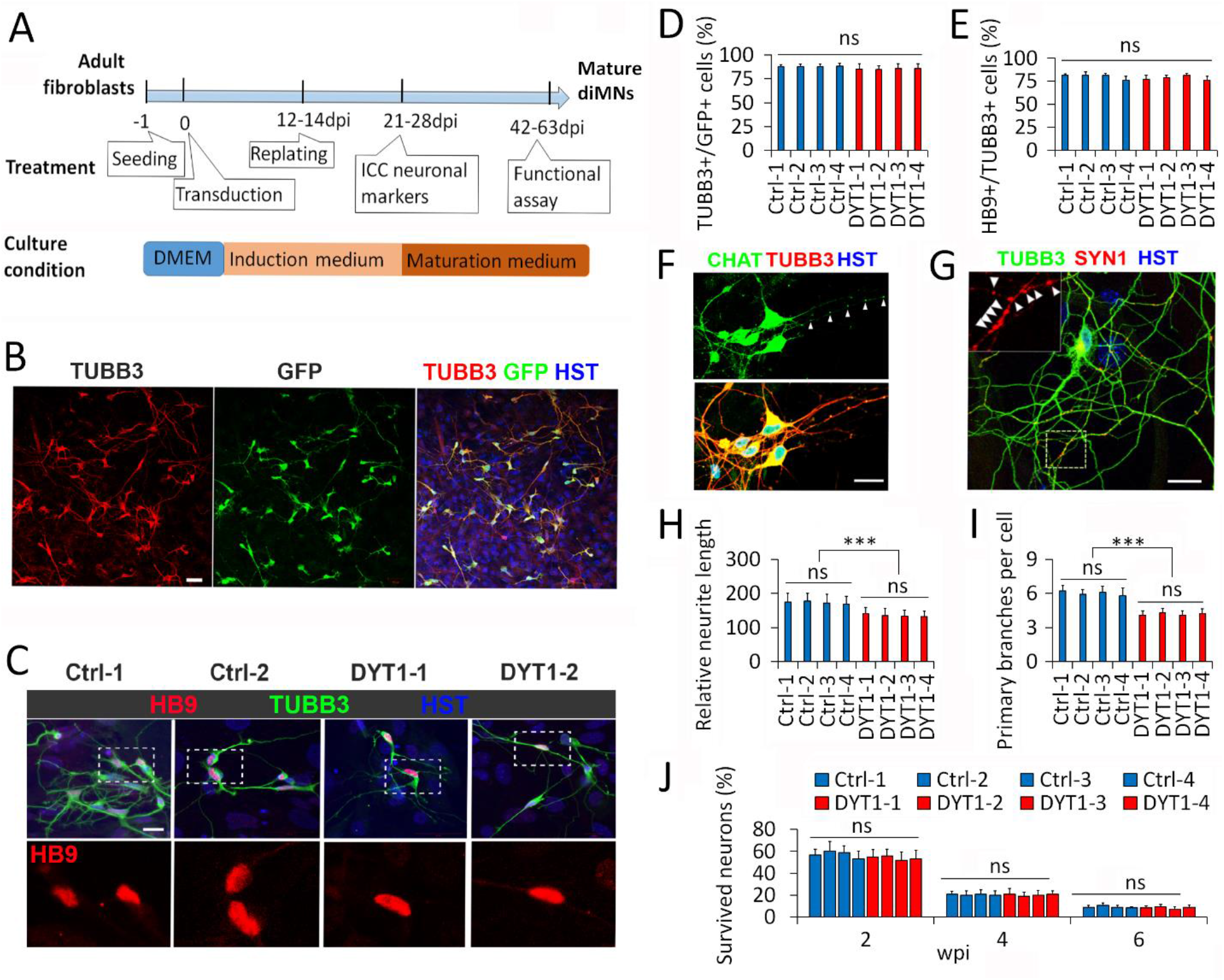
Direct conversion of DYT1 patient fibroblasts to motor neurons. **(A)** A schematic of the direct conversion procedure. **(B)** A lower magnification view of the reprogrammed neurons at 19 days post viral infection (dpi). Virus-transduced cells are indicated by the co-expressed GFP. Scale bar: 50 μm. **(C)** Confocal images of the reprogrammed neurons at 19 dpi. Higher magnification views of nuclear HB9 are also shown. Scale bar: 20 μm. **(D)** Quantification of the reprogramming efficiency (n = 259 for Ctrl-1, n = 249 for Ctrl-2, n = 237 for Ctrl-3, n = 243 for Ctrl-4, n = 257 for DYT1-1, n = 254 for DYT1-2, n = 302 for DYT1-3, and n = 257 for DYT1-4 from triplicates; ns, not significant). **(E)** Fractions of diMNs among the reprogrammed neurons (n = 257 for Ctrl-1, n = 273 for Ctrl-2, n=252 for Ctrl-3, n = 281 for Ctrl-4, n = 229 for DYT1-1, n = 219 for DYT1-2, n = 289 for DYT1-3, and n = 233 for DYT1-4 from triplicates; ns, not significant). **(F)** Robust expression of the motor neuron marker CHAT at 7 wpi. Arrowheads indicate CHAT+ puncta. Scale bar: 20 μm. **(G)** Expression of the presynaptic marker SYN1 at 9 wpi. The inset shows discrete SYN1+ puncta (indicated by arrow heads). Scale bar: 50 μm. **(H)** Quantitative analysis of neurite length at 2 wpi (n = 123 for Ctrl-1, n = 126 for Ctrl-2, n = 137 for Ctrl-3, n = 125 for Ctrl-4, n = 114 for DYT1-1, n = 127 for DYT1-2, n = 118 for DYT1-3, n = 113 for DYT1-4 from triplicates; ns, not significant; ****p<0.0001). **(I)** Neurite number of the primary branches examined at 4 wpi (n = 163 for Ctrl-1, n = 153 for Ctrl-2, n= 157 for Ctrl-3, n = 145 for Ctrl-4, n = 152 for DYT1-1, and n = 143 for DYT1-2, n = 125 for DYT1-3, and n = 131 for DYT1-4 from triplicates; ns, not significant; *** p<0.001). **(J)** A time-course analysis of diMN survival (each line n (neurons) > 2000 from triplicates; ns, not significant).

### DYT1 diMNs Show Impaired Development but Normal Survival

DYT1 dystonia can also be classified as a neurodevelopmental disorder ^3^. To track the development of human DYT1 diMNs, we examined neurite outgrowth and primary branches at the early reprogramming stages of 2 and 4 wpi as described in previous studies ^27–30^. DYT1 diMNs had significantly shorter neurites (Fig. 1H) and many fewer primary branches (Fig. 1I) than controls. On the other hand, no significant differences were identified on neuronal survival within 6 weeks (Fig. 1J). These results support the notion that DYT1 dystonia is caused by alterations in neuronal function rather than by neurodegeneration ^31^.

### DYT1 diMNs Exhibit Dysfunctional Nucleocytoplasmic Transport (NCT)

Torsin has been shown to participate in nuclear export of mRNAs via a non-conventional mechanism called nuclear envelope budding in *Drosophila* ^32^. In torsin mutants, disruption of mRNA nuclear export led to slowed growth and impaired development and plasticity of neuromuscular junctions ^14^. To examine whether mRNA export was similarly impaired in DYT1 diMNs, we performed fluorescence *in situ* hybridization (FISH), as previously described ^25,33,34^. We used digoxigenin-labeled oligo-dT as probes to detect the subcellular distribution of Poly (A) mRNAs. FISH specificity was confirmed by a lack of signals when samples were treated with RNaseA or when digoxigenin-labeled oligo-dA was used as a probe (Fig. 2A, B). mRNA nuclear export was measured by the ratio of nuclear oligo-dT signals to cytoplasmic signals (dT_nuc_/dT_cyt_). In healthy control diMNs, a majority of mRNAs was localized in the cytoplasm when examined at 6 wpi (Fig. 2C, D). In sharp contrast, substantial amounts of mRNAs were accumulated inside the nucleus of DYT1 diMNs (Fig. 2C, D).

**Figure 2.**
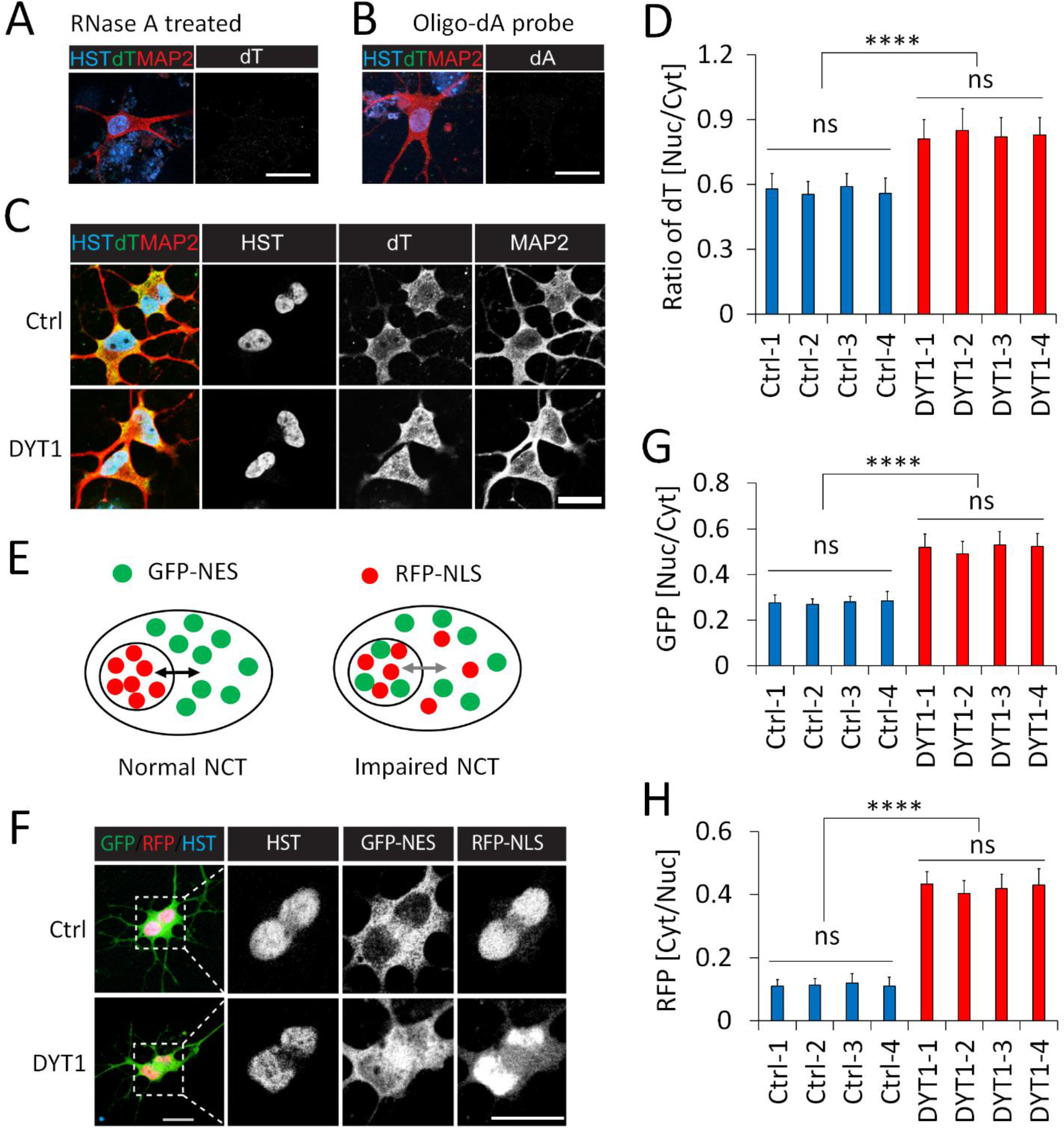
Impaired nucleocytoplasmic transport in DYT1 diMNs. **(A)** RNase A-treated diMNs as negative controls for fluorescence in situ hybridization (FISH). Cells were analyzed at 6 wpi. Scale bar: 20 μm. **(B)** Oligo-dA probes as negative controls for FISH assays at 6 wpi. Scale bar: 20 μm. **(C)** FISH analysis of diMNs with oligo-dT probes at 6 wpi. Scale bar: 20 μm. **(D)** Quantification of oligo-dT signal distribution (n = 65 for Ctrl-1, n = 67 for Ctrl-2, n = 72 for Ctrl-3, n= 70 for Ctrl-4, n = 64 for DYT1-1, n = 63 for DYT1-2, n = 74 for DYT1-3, and n = 68 for DYT1-4 from four replicates; ns, not significant; ****p<0.0001). **(E)** A dual reporter system for measuring protein nucleocytoplasmic transport (NCT). NES: nuclear export signal; NLS: nuclear localization signal. **(F)** Representative confocal images of reporter distribution in diMNs at 6 wpi. Scale bar: 20 μm. **(G)** Subcellular distribution of GFP-NES (n = 62 for Ctrl-1, n = 64 for Ctrl-2, n = 76 for Ctrl-3, n = 74 for Ctrl-4, n = 67 for DYT1-1, and n = 68 for DYT1-2 n = 75 for DYT1-3, and n = 73 for DYT1-4 from four replicates; ns, not significant; ****p<0.0001). Subcellular distribution of RFP-NLS (n = 62 for Ctrl-1, n = 64 for Ctrl-2, n = 76 for Ctrl-3, n = 74 for Ctrl-4, n = 67 for DYT1-1, and n = 68 for DYT1-2 n = 75 for DYT1-3, and n = 73 for DYT1-4 from four replicates; ns, not significant; ****p<0.0001). **(H)** Subcellular distribution of RFP-NLS (n = 34 for WT and n = 36 for DYT1 from triplicates; **p<0.01).

Interestingly, we noticed obvious MAP2 signals in the nucleus of DYT1 diMNs (Fig. 2C), suggesting that protein NCT might also be compromised. We used a dual reporter system (2Gi2R) to further examine NCT ^25^. This reporter system consists of a fused double-GFP with a nuclear export signal (2GFP-NES) and a double RFP containing a nuclear localization signal (2RFP-NLS) ^35^. In cells with normal NCT activity, GFP and RFP are localized in the cytoplasm and nucleus, respectively; in cells with compromised NCT, this subcellular distribution will be disrupted (Fig. 2E). An increased ratio of nuclear to cytoplasmic GFP (GFP_nuc_/GFP_cyt_) represents impaired protein export, while a higher ratio of cytoplasmic to nuclear RFP (RFP_cyt_/RFP_nuc_) indicates compromised protein nuclear import. Both GFP and RFP were mainly localized in the cytoplasm and nucleus, respectively, in control diMNs when examined at 6 wpi (Fig. 2F-H). In contrast, DYT1 diMNs had a substantial amount of GFP accumulated inside the nucleus and obvious RFP detected in the cytoplasm (Fig. 2F). As such, the ratios of GFP_nuc_/GFP_cyt_ and RFP_cyt_/RFP_nuc_ were significantly higher for DYT1 diMNs than control neurons (Fig. 2G and H). Together, FISH analysis and dual reporter assays clearly indicate that both mRNA nuclear export and protein NCT are impaired in DYT1 diMNs.

### NCT Impairments in DYT1 iPSC-MNs

To verify our findings, we generated MNs via iPSC-based reprogramming and differentiation (Fig. 3A). The fibroblasts from a healthy control and a patient with GAG deletion in *TOR1A* gene were reprogrammed into iPSCs (Fig. S2), which were then differentiated into neural progenitors ^36^ (Fig. S3). Neural progenitors were further induced to become MNs (Fig. S4), as previously reported ^24^. Markers for mature neurons, such as CHAT (Fig. S4B), MAP2 (Fig. S4E), and presynaptic marker SYN1 (Fig. S4F), were detectable within 3 wpi in these iPSC-MNs. Electrophysiological properties were examined by whole-cell patch-clamp recordings at 4 wpi. All patched iPSC-MNs fired repetitive action potentials upon current injections (Fig. S5A). No apparent differences were observed between DYT1 and control iPSC-MNs on action potential frequency, delay of the first spike, and rise and decay of Vmax (Fig. S5B-E).

**Fig. 3.**
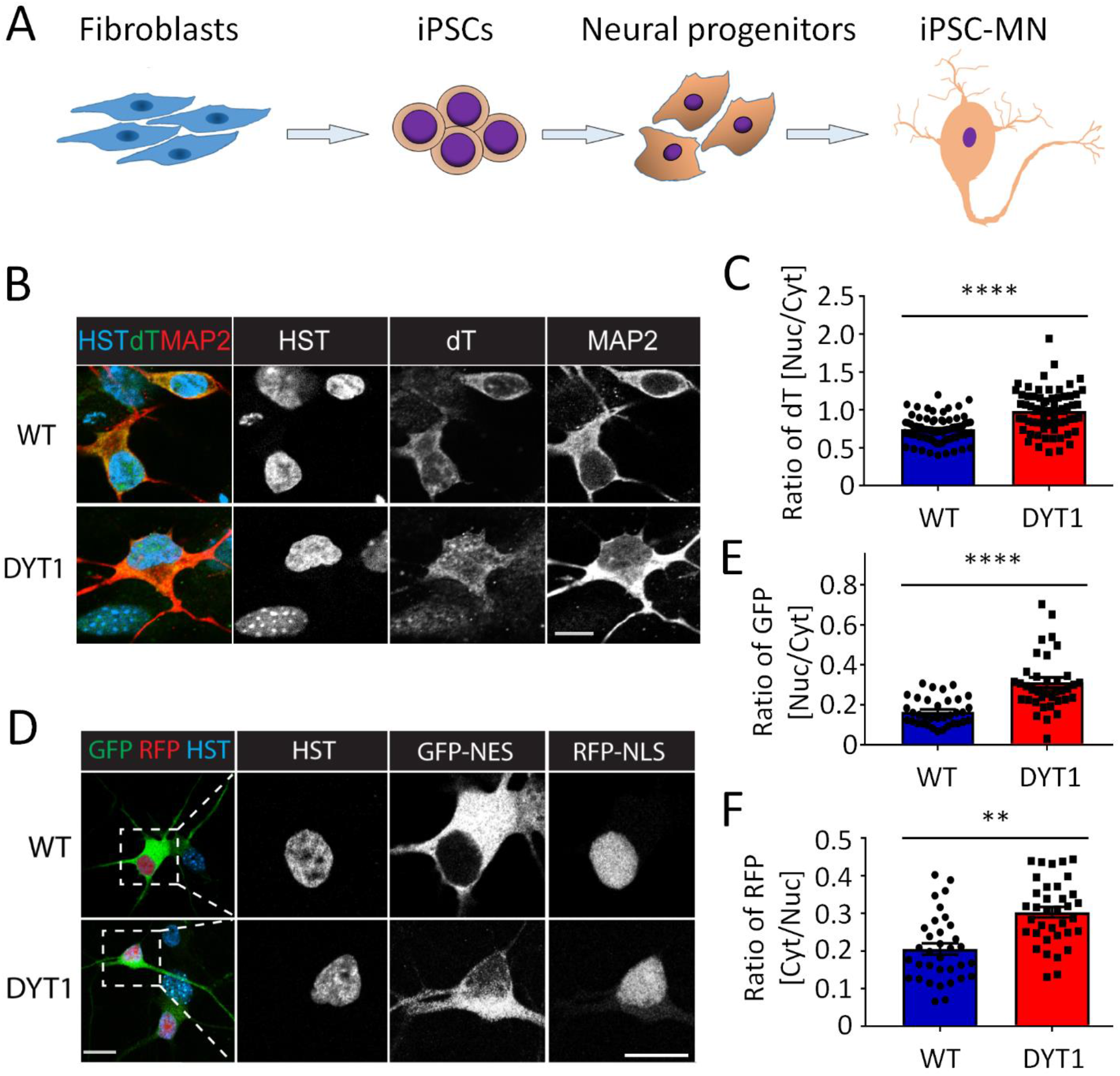
NCT impairments in DYT1 iPSC-MNs. **(A)** A schematic of the iPSC-based procedure to generate MNs from adult fibroblasts. **(B)** Representative confocal images of FISH assay in iPSC-MNs at 3 wpi. Scale bar: 10 μm. **(C)** Subcellular distribution of mRNAs by FISH assay (n= 85 for WT and 73 for DYT1 from triplicates; ****p<0.0001). **(D)** Representative confocal images of iPSC-MNs with the dual reporter at 3 wpi. Scale bar: 20 μm. **(E)** Subcellular distribution of GFP-NES (n = 34 for WT and n = 36 for DYT1 from triplicates; ****p<0.0001).

mRNA nuclear export and protein nucleocytoplasmic transport were also significantly compromised in DYT1 iPSC-MNs (Fig. 3B-F), though these defects were less severe than in DYT1 diMNs (Fig. 2). For example, the cytoplasmic localization of RFP-NLS was less obvious in DYT1 iPSC-MNs than DYT1-diMNs (Fig. 3D compared to Fig. 2F). Such a difference may reflect an aging effect on disease phenotypes, since diMNs retain aging-associated features whereas iPSC-MNs are rejuvenated during the reprogramming process ^24,35^.

### Abnormal Nuclear Envelope but no Perinuclear “Blebs” in DYT1 MNs

One cellular hallmark in *Tor1a* mutant mice (*Tor1a* knockout or *Tor1a*^*ΔE/ΔE*^) is the abnormal NE morphology in multiple areas of the CNS ^10,37^. It is characterized by an accumulation of “blebs” in the perinuclear space, morphologically similar to the budding granules in the body-wall muscle in torsin-mutant flies ^14,32^. We first examined whether such abnormalities also occur and contribute to impaired NCT in human DYT1 neurons. Nuclear morphology was studied by transmission electron microscopy (TEM). At the iPSC stage, both WT and DYT1 cells had smooth and round nuclei with clear double nuclear membranes, except for a few small wrinkles in some DYT1 iPSCs (Fig. 4A and B). After differentiation into mature MNs, most WT cells retained the regular, smooth nuclear shapes with occasional wrinkles and light invaginations (Fig. 4C). In contrast, the nuclear morphology of DYT1 iPSC-MNs showed many sharp angles and deep invaginations (Fig. 4D-H). However, we failed to observe any “bleb” structures at the perinuclear space in DYT1-MNs from three batches of samples. If “bleb” structures exist, the frequency must be very low. Such a result indicates that the mechanisms underlying the abnormal NCT might differ between human DYT1 patients and other animal models such as *Drosophila* ^14^ and mouse ^10,37^.

**Fig. 4.**
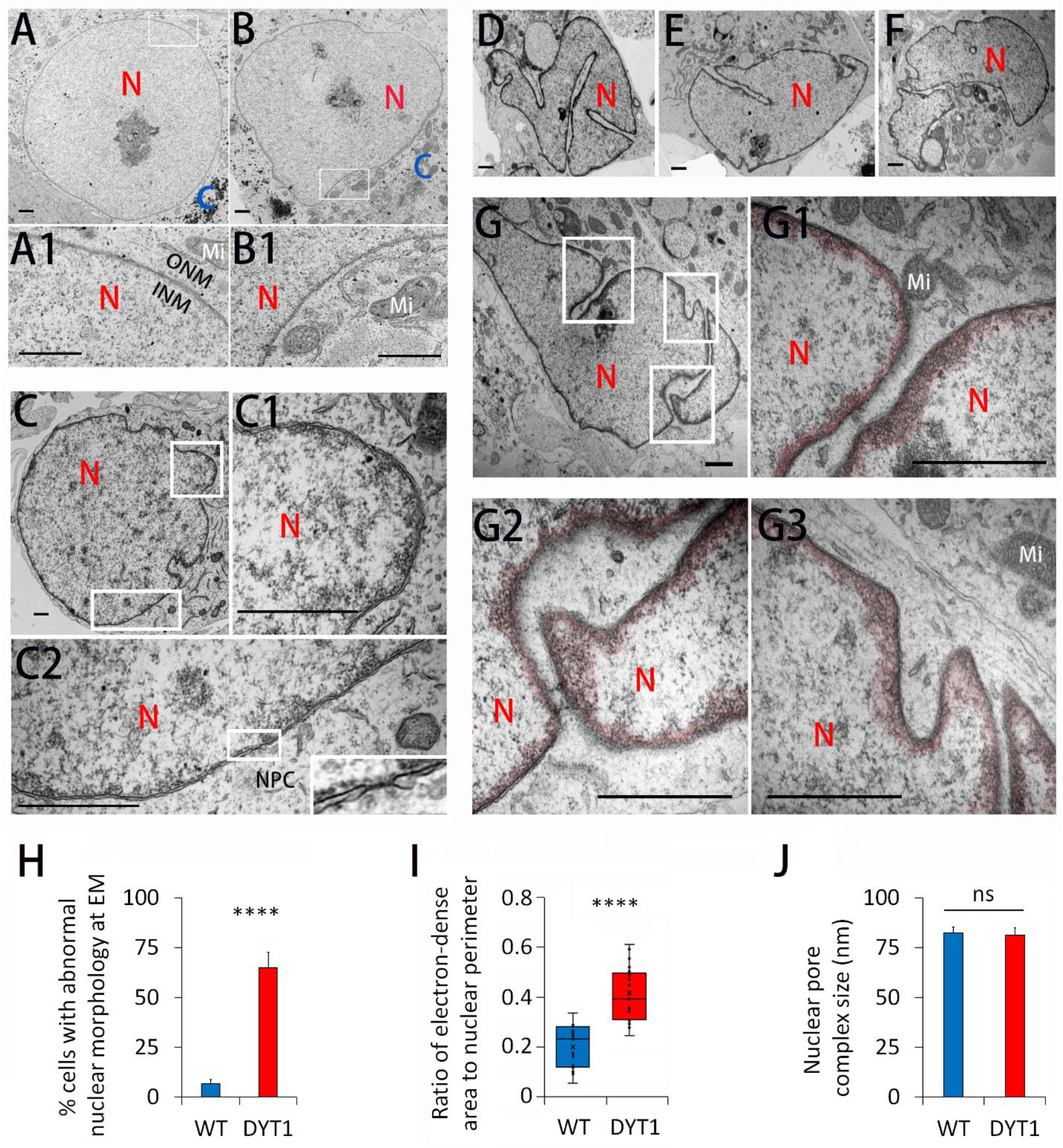
Nuclear envelope deformation in DYT1 MNs. **(A)** Representative electron micrograph of WT iPSC. A higher magnification view of the boxed region is shown in A1 at the bottom panel. N, nucleus; C, cytoplasm; INM, inner nuclear membrane; ONM, outer nuclear membrane; Mi, mitochondria. Scale bars: 1 μm. **(B)** Representative electron micrograph of DYT1 iPSC. A higher magnification view of the boxed region is shown in B1 at the bottom panel. Scale bars: 1 μm. **(C)** Representative electron micrographs of WT iPSC-MN at 3 wpi. Higher magnification views of the boxed regions are shown in C1 and C2, respectively. NPC, nuclear pore complex. Scale bars: 1 μm. **(D-G)** Representative electron micrographs of DYT1 iPSC-MNs at 3 wpi. Higher magnification views of the boxed regions are shown in G1, G2 and G3, respectively. The areas of electronic dense beneath the nuclear membrane are highlighted in red. Scale bars: 1 μm. **(H)** iPSC-MNs with abnormal nuclear morphology at 3 wpi (n (nuclei) = 47 for WT and 42 for DYT1 from triplicates; ****<0.0001). **(I)** Ratio of electron dense area immediately beneath the nuclear membrane to the nuclear perimeter in iPSC-MNs at 3 wpi (n (nuclei) = 24 for WT and 22 for DYT1 from triplicates; ****<0.0001.) (**J**) Nuclear pore size in iPSC-MNs at 3 wpi (n (nuclei/nuclear pore) = 17/56 for WT and 21/34 for DYT1 from triplicates; ns, not significant).

Interestingly, we found that DYT1 MNs exhibited thickened nuclear lamina with high electron-dense areas immediately beneath the nuclear membranes, especially in regions with abnormal angles and invaginations (Fig. 4G1-G3). Quantification showed a much higher ratio of the electron-dense area to the nuclear perimeter in DYT1-MNs than controls (Fig. 4I). On the other hand, the nuclear pore complex (NPC) size was comparable between WT- and DYT1-MNs (Fig. 4J). We further examined several nuclear pore subunits by immunocytochemistry and confocal microscopy. The signal density of Nup98, Nup358, or other nucleaporins (Mab414) was similar among the examined samples (Fig. S6). These results indicate that the impaired NCT in DYT1 neurons might not be due to NPC malformation.

### LMNB1 Mislocalization in DYT1 MNs

Since thickening of the nuclear lamina is a salient feature of DYT1 MNs (Fig. 4G and I), it was further examined by immunocytochemistry and confocal microscopy. Nuclear lamina consists of A-type and B-type lamins, which are intermediate filament proteins and form the meshwork beneath the inner nuclear membrane to support the nuclear morphology and structural integrity of the nuclear envelope ^38–41^. Lamin A (LMNA) and lamin C (LMNC) are the major A-type lamins encoded by a single *LMNA* gene in humans ^39^. Staining for LMNA/C showed their exclusive localization in the nuclear periphery in both control and DYT1 diMNs (Fig. 5A). However, unlike the regular ovoid nucleus with a smooth outline in control cells, approximately 60% of DYT1 diMNs showed abnormal nuclear morphology with sharper and rigid angles (Fig. 5A and B), consistent with what was observed by TEM (Fig. 4D-H).

**Fig. 5.**
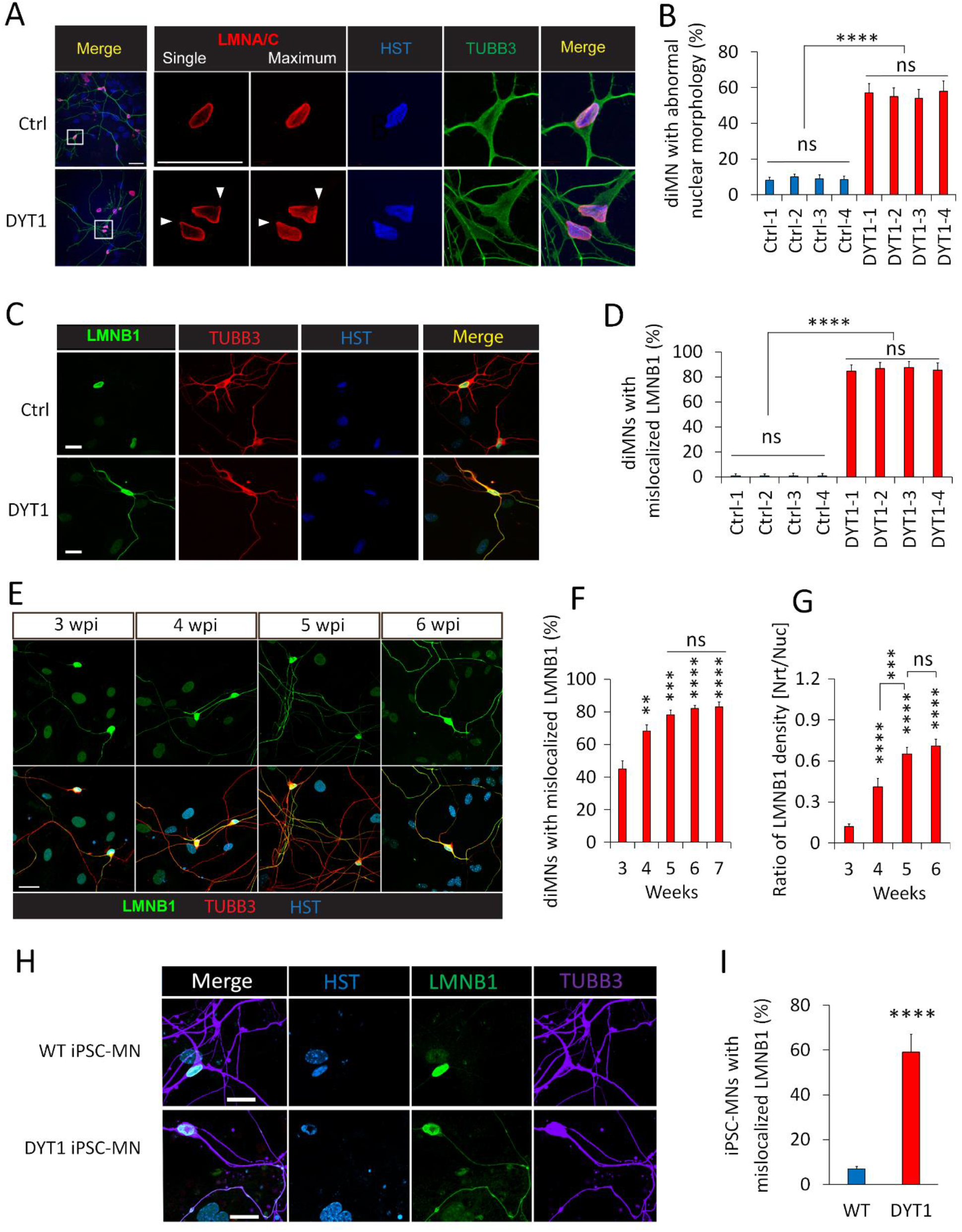
Abnormal nuclear morphology and mislocalized LMNB1 in DYT1 diMNs. **(A)** Representative confocal images of diMNs at 4 wpi. Arrowheads indicate the sharp and rigid angles in nuclear membrane. Scale bar: 50 μm. **(B)** Quantification of diMNs with abnormal nuclear morphology at 4 wpi (n > 100 for each line from four replicates; ns, not significant; ****p<0.0001). **(C)** Representative confocal images of diMNs at 6 wpi. Scale bar: 20 μm. **(D)** Percentages of diMNs with mislocalized LMNB1 at 6 wpi (n = 410 for Ctrl-1, n = 418 for Ctrl-2, n= 490 for Ctrl-3, n = 378 for Ctrl-4, n = 358 for DYT1-1, n = 349 for DYT1-2, n = 378 for DYT1-3, and n = 376 for DYT1-4 from four replicates; ns, not significant; **** p<0.0001). **(E)** A time course analysis of LMNB1 mislocalization in DYT1 diMNs. Scale bar: 50 μm. **(F)** Frequency of LMNB1 mislocalization in DYT1 diMNs at the indicated time points (n = 458 for 3wpi, n= 425 for 4 wpi, n = 327 for 5 wpi, n = 314 for 6 wpi, and n= 213 for 7 wpi from triplicates; ns, not significant; compare to 3 wpi, ** p<0.01, *** p<0.001, **** p<0.0001). **(G)** Subcellular distribution of LMNB1 (neurite vs. nucleus) in DYT1 diMNs at the indicated time points (n = 126 for 3 wpi, n= 114 for 4 wpi, n = 128 for 5 wpi and, n = 107 for 6 wpi from triplicates; ns, not significant; *** p<0.001, ****p<0.0001). **(H)** Representative confocal images of iPSC-MNs at 3 wpi. Scale bar: 20 μm. **(I)** Subcellular localization of LMNB1 in iPSC-MNs at 3 wpi. (n = 397 for WT and n = 423 for DYT1 from triplicates; ****p<0.0001).

Similar to LMNA/C, staining of the major B-type nuclear lamins LMNB1 revealed its normal nuclear localization in control diMNs (Fig. 5C). Unexpectedly, a substantial amount of LMNB1 were detected in the cytoplasm and neurites of DYT1 diMNs when examined at 6 wpi (Fig. 5C). Quantification showed approximately 80% of DYT1 diMNs had mislocalized LMNB1, whereas such mislocalization was rarely detected in healthy controls (Fig. 5D). Co-staining with SMI-32 and MAP2 indicated that the mislocalized LMNB1 was distributed in both axons and dendrites (Fig. S7). A time-course analysis revealed that cytoplasmic LMNB1 in DYT1 diMNs became progressively more frequent, with 45% at 3 wpi and 80% at 6-7 wpi (Fig. 5E-F). The cytoplasmic fraction of LMNB1 also greatly increased with time and reached a plateau at 6 wpi (Fig. 5G). Importantly, LMNB1 mislocalization was also observed in DYT1 iPSC-MNs (Fig. 5H). Such mislocalization could be detected as early as 1.5 wpi and in about 60% of DYT1 iPSC-MNs when examined from 2 to 8 wpi (Fig. 5I). In contrast, we failed to detect any obvious mislocalization of several other nuclear proteins, including LMNA/C (Fig. 5A) and the nuclear pore subunits NUP98 and NUP358 in DYT1 diMNs (Fig. S6).

### TOR1A Hypoactivity Disrupts NCT and LMNB1 Localization

To determine whether these above defects were caused by dysregulation of TOR1A or largely influenced by the genetic background of human patients, we employed two approaches. One was to ectopically express either TOR1A or TOR1AΔE mutant in WT iPSC-MNs. These MNs could be identified by the co-expressed GFP reporter. Western blot analysis showed that both TOR1A and TOR1AΔE were expressed at a similar level (Fig. 6A). Interestingly, ectopic TOR1AΔE mutant but not the wild type TOR1A disrupted mRNA nuclear export (Fig. 6B) and LMNB1 subcellular distribution (Figs. 6C and D). In a second approach, we downregulated *TOR1A* expression through shRNAs in WT iPSC-MNs. shRNA-expressing cells were marked by the mCherry reporter. Each of the two separate shRNAs against *TOR1A* could efficiently down-regulate its expression at both protein and mRNA levels (Figs. 6E and F). Consistently, downregulation of endogenous *TOR1A* led to defective export of nuclear mRNAs (Fig. 6G) and altered subcellular localization of LMNB1 (Fig. 6H). This LMNB1 mislocalization was independent of the neuronal differentiation process because MNs infected with the *TOR1A-shRNA* virus at either pre-(neuronal progenitors) or post-differentiation (MNs at 5 dpi) stages exhibited similar results (Fig. 6I). Together, these results strongly support that NCT impairment and LMNB1 mislocalization are caused by TOR1A hypoactivity.

**Fig. 6.**
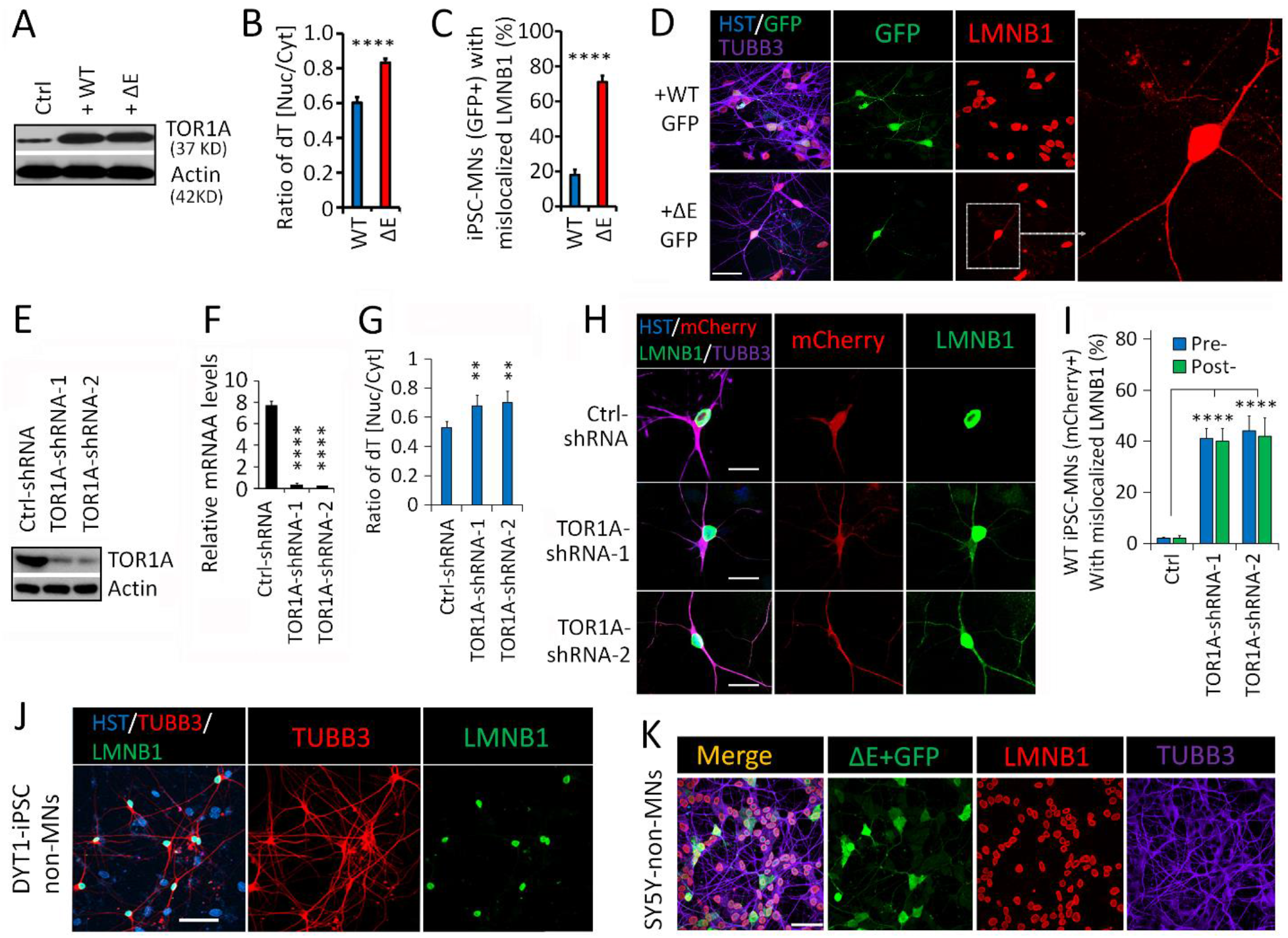
TOR1A hypoactivity disrupts NCT and LMNB1 localization in a MN-specific manner. **(A)** Protein expression of the ectopically expressed TOR1A WT or ΔE in iPSC-MNs; Actin serves as a loading control. **(B)** The effect of TOR1A expression on mRNA distribution assayed by FISH in iPSC-MNs at 3 wpi (n = 99 for WT and n = 78 for DYT1 from triplicates; ****p<0.0001). **(C)** Quantification of LMNB1 mislocalization in iPSC-MNs at 3 wpi (n = 82 for WT and n = 75 for DYT1 from triplicates; ****p<0.0001). **(D)** Representative confocal images of iPSC-MNs with ectopic (GFP+) TOR1A WT or ΔE at 3 wpi. Scale bar: 50 μm. **(E)** Western blots showing shRNA knockdown of TOR1A protein expression in iPSC-MNs. **(F)** qRT-PCR showing downregulation of *TOR1A* mRNA by shRNAs (n = 3; ****<0.0001). **(I)** The effect of *TOR1A* downregulation on mRNA subcellular distribution assayed by FISH in iPSC-MNs at 3 wpi (n =95 for Ctrl, n = 87 for TOR1A-shRNA-1 and n = 78 for TOR1A-shRNA-2 from triplicates; **<0.01). **(J)** Representative confocal images of iPSC-MNs with the indicated shRNAs (mCherry+) at 3 wpi. Scale bar: 25 μm. The effect of *TOR1A* downregulation on LMNB1 subcellular distribution in iPSC-MNs. *ShRNA* lentiviruses were added at either pre-(neural progenitors) or post-differentiation (MN at 5dpi) stages. n (pre- and post-) = 126 and 113 for Ctrl, n = 118 and 97 for *TOR1A-shRNA-1*, and n = 103 and 94 for *TOR1A-shRNA-2* from triplicates; ***p<0.001. **(K)** Representative confocal images of DYT1-iPSC spontaneously differentiated non-MNs at 3 wpi. Scale bar: 100 μm. Representative confocal images of SY5Y-derived non-MNs with ectopic expression of TOR1AΔE (GFP+) at 4 wpi. Scale bar: 50 μm.

To examine whether hypoactivity of TOR1A also leads to LMNB1 mislocalization in other neuron subtypes, we generated non-MNs via two approaches. One was spontaneous differentiation of DYT1 neural progenitors into generic neurons (Fig. 6J), which mainly consisted of glutamatergic and GABAergic neurons ^24^. The second approach was to ectopically express *TOR1AΔE* in generic neurons that were differentiated from human SH-SY5Y neuroblasts (Fig. 6K). Interestingly, we did not notice any mislocalized LMNB1 in either of these generic neurons (Fig. 6J and K). These results suggest that TOR1A hypoactivity may cause LMNB1 mislocalization in a MN-specific manner.

### Specific Upregulation of LMNB1 in DYT1 MNs

LMNB1 expression in iPSC-MNs was then examined by western blot analysis. Beta-actin and neuron-specific microtubules TUBB3 were used as protein loading controls. Strikingly, the LMNB1 protein level was dramatically higher (2.5 folds) in DYT1 iPSC-MNs than in WT controls (Fig. 7A and B). Such a result was consistent with immunocytochemistry showing brighter LMNB1 signals and cytoplasmic accumulation in DYT1 MNs (Figs. 5C). Notably, qRT-PCR analysis showed that *LMNB1* transcripts in DYT1 iPSC-MNs were 3-5 fold higher than those in WT controls (Fig. 7C), indicating that increased LMNB1 protein was mainly contributed by upregulated gene expression. In sharp contrast, no significant differences were observed for lamin*-* associated proteins, LAP2A and LAP2B (Fig. 7A and B), which are known to interact with LMNA/C and LMNB1 ^42^. We also failed to detect any significant changes on LMNB2 (another B-type nuclear lamina), NUP98 and NUP358 (subunits of the NPCs), GLE1 (a factor involved in mRNA export), and RANGAP1 (a nuclear pore-associated protein involving in nuclear transport) (Fig. 7A and B). Interestingly, LMNB1 protein levels were also increased in WT-MNs when *TOR1A* was downregulated by shRNAs (Fig. 7D). These results indicate that LMNB1 is uniquely dysregulated on both protein expression and subcellular localization in DYT1 MNs or MNs with hypoactive TOR1A.

**Fig. 7.**
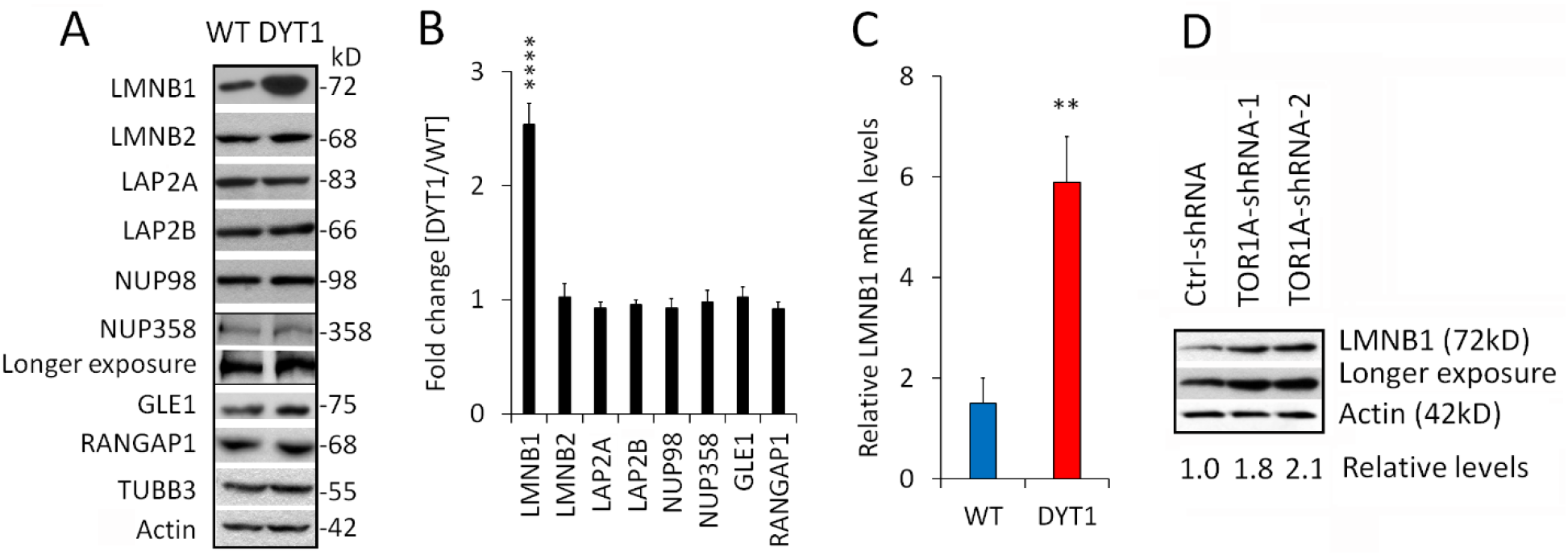
Specific upregulation of LMNB1 in DYT1 MNs. **(A)** Western blots for the indicated proteins in WT-and DYT1 iPSC-MNs at 11 dpi. TUBB3 and Actin are loading controls. Both shorter and longer exposure of NUP358 are also shown. **(B)** Fold changes (DYT1 to WT) of the indicated proteins in (A) after normalization to the loading control (n=3). **(C)** Relative *LMNB1* mRNA levels analyzed by qRT-PCR in iPSC-MNs at 11 dpi (n = 3). **(D)** Western blots for LMNB1 expression (shorter and longer exposure) in iPSC-MNs with the indicated shRNAs at 11 wpi. Actin serves as a loading control. Values at the bottom indicate the relative LMNB1 protein levels.

### Phenotypic Amelioration of DYT1 MNs via LMNB1 Downregulation

To determine whether LMNB1 dysregulation contributes to the pathological phenotypes in DYT1 MNs, we knocked down its expression through a shRNA approach. The shRNA-expressing cells were marked with the co-expressed mCherry reporter. A shRNA against luciferase was used as a control. Two shRNAs were identified to be efficient in reducing the expression of endogenous LMNB1 determined by both western blots and immunocytochemistry (Fig. 8A and B). Interestingly, downregulation of LMNB1 resulted in a much reduced frequency of LMNB1 mislocalization in DYT1 MNs (Fig. 8C). It also significantly improved neurite outgrowth (Fig. 8D), increased the number of primary branches (Fig. 8E), and markedly reduced the percentage of cells with abnormal nuclear morphology (Fig. 8F). When compared to the control shRNA, FISH analysis with digoxigenin-labeled oligo-dT probes showed that *LMNB1*-shRNAs significantly reduced nuclear accumulation of mRNAs in DYT1-MNs (Figs. 8G and H). Together, these results clearly indicate that LMNB1 dysregulation is a salient pathological feature contributing to the delayed neuron development, abnormal nuclear morphology, and impaired nucleocytoplasmic transport.

**Fig. 8.**
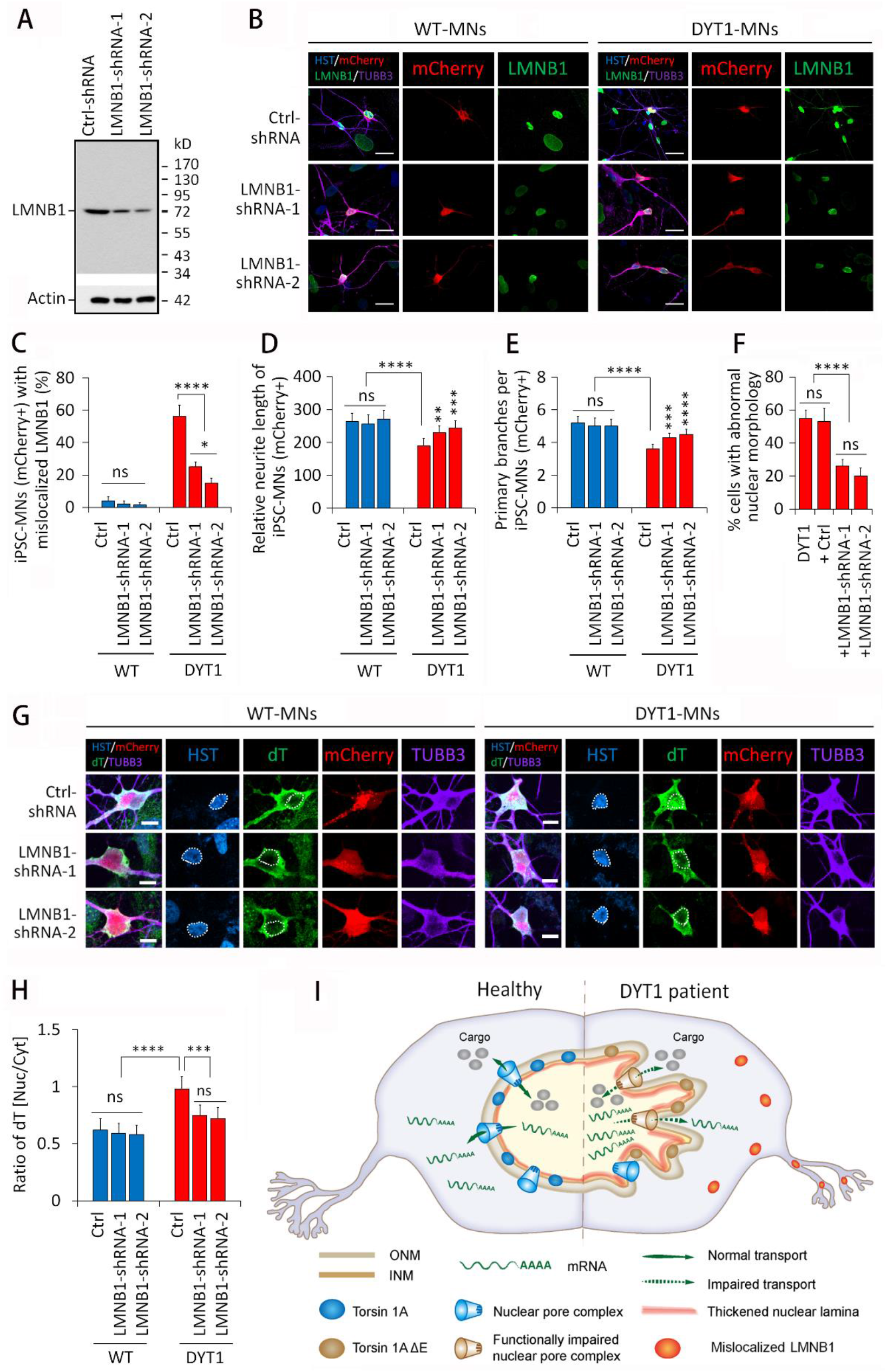
Phenotypic amelioration of DYT1 MNs via LMNB1 downregulation. **(A)** Western blots showing shRNA-mediated downregulation of LMNB1. The single band of LMNB1 on the whole blotting membrane indicates the specificity of LMNB1 antibody. Actin serves as a loading control. **(B)** Representative confocal images of iPSC-MNs with the indicated shRNAs (mCherry+) at 3 wpi. Scale bar: 25 μm. **(C)** Percentage of iPSC-MNs (mCherry+) with mislocalized LMNB1 in (B). (n (neurons) > 200 for each condition from triplicates; ns, not significant; * p<0.05, ****p<0.0001). **(D)** Quantitative analysis of neurite length of iPSC-MNs (mCherry+) at 2 wpi under the indicated conditions (n (neurons) > 100 for each sample from triplicates; ns, not significant; ** p<0.01, *** p<0.001, ****p<0.0001). **(E)** Neurite number of the primary branches in iPSC-MNs (mCherry+) at 4 wpi. (n (neurons) > 100 each sample from triplicates; ns, not significant; *** p<0.001, ****p<0.0001). **(F)** Percentage of DYT1 iPSC-MNs with abnormal nuclear morphology at 3 wpi (n = 104 for DYT1, n = 98 for Ctrl-shNRA, n = 72 for LMNB1-shRNA1 and n = 69 for LMNB1-shNRA2 from triplicates; ns, not significant; ****p<0.0001). **(G)** Representative confocal images of FISH assay of iPSC-MNs at 3 wpi. *shRNA*-expressing cells are identified by the co-expressed mCherry. Dotted lines highlight nuclei. Scale bar: 10 μm. **(H)** Quantification of oligo-dT signal distribution in (G). (n (neurons) > 100 for each sample from triplicates; ns, not significant; ***p<0.001, ****p<0.0001). **(I)** A working model for DYT1 pathogenesis. The healthy MNs have normal nuclear morphology and NCT activity. In DYT1 MNs, LMNB1 upregulation leads to thickened nuclear lamina, cytoplasmic localization, abnormal nuclear morphology, and impaired NCT, which may subsequently lead to mislocalization of mRNAs and proteins and dysfunction of neurons.

## Discussion

### Reprogrammed Human Neurons as Novel Models for DYT1 Dystonia

Most cases of DYT1 dystonia are caused by a single amino-acid deletion in TOR1A under a heterozygous background. Interestingly, mice with the identical *Tor1a* mutation as a heterozygote failed to show any pathological phenotypes ^10^, suggesting that species-specific differences may play a role in DYT1 pathology. Nonetheless, the limited access to patient neurons has greatly impeded the progress of DYT1 dystonia research. In this study, we generated DYT1 MNs via two strategies: direct conversion from patient fibroblasts and iPSC-based reprogramming and differentiation. These neurons are in the same genetic background as human patients and maintain endogenous levels of TOR1A expression. As such, they provide an unprecedented *in vitro* cellular model for dystonia research.

DYT1 MNs from both strategies show similar phenotypes including delayed neuron development, abnormal nuclear morphology, and disrupted mRNA nuclear export and protein nuclear transport (Table S2). Unlike iPSCs, which are epigenetically reset to an embryonic state ^43,44^, directly reprogramed neurons from donors retain features associated with age and disease ^24,35^. We consistently find that DYT1 diMNs exhibit more severe defects than DYT1 iPSC-MNs in neurite outgrowth, nuclear morphology, and NCT (Table S2). However, the iPSC-based strategy has several unique advantages: both the yield and the purity of iPSC-MNs are much higher than diMNs, and the culture time to obtain mature neurons is also shorter (Table S2). Therefore, iPSC-MNs are suitable for experiments requiring a large amount of samples, whereas diMNs can provide aging-relevant neurons for single cell analysis. Both strategies should be valuable in DYT1 dystonia research.

Our study further provides evidence that lower motor neurons are implicated in the pathogenesis of dystonia. We reveal for the first time that LMNB1 is upregulated and mislocalized in DYT1 MNs. Most interestingly, downregulation of LMNB1 can significantly improve development, NE morphology, and mRNA nuclear export of DYT1 MNs. Our results support a model that upregulated LMNB1 causes thickening of the nuclear lamina, thereby leading to abnormal nuclear morphology and impaired nucleocytoplasmic transport in DYT1 MNs (Fig. 8I). Progressive LMNB1 accumulation and subcellular mislocalization are key pathological features of DYT1 MNs and may constitute a novel molecular target for intervention.

### Increased Nuclear LMNB1 Causes Abnormal NE Morphology

Nuclear LMNB1 is an essential scaffolding component of the lamina meshwork beneath the nuclear envelope. It plays important roles in determining nuclear morphology, DNA regulation, and gene expression ^45,46^. Overexpression of LMNB1 is shown to increase nuclear rigidity ^47^. This is consistent with our result that LMNB1 overexpression can disrupt nuclear morphology in fibroblasts (Fig. S8). An abnormal extra copy of *LMNB1* gene causes autosomal dominant leukodystrophy (ADLD) characterized by loss of myelin in the brain and the spinal cord, leading to movement problems ^48–50^. Consistently, we find that LMNB1 expression is obviously increased in DYT1 MNs, and the NE morphology is abnormal at both light- and electron-microscope levels. Downregulation of LMNB1 in DYT1 MNs can alleviate the defects in neuron development, NE morphology, and mRNA nuclear export, suggesting that increased LMNB1 contributes to these abnormalities in DYT1 neurons.

Despite intensive efforts examining the nuclear envelope structure by TEM, we failed to detect bleb-like structures in DYT1 MNs. Such a result is in sharp contrast to what was observed in *Drosophila* with torsin mutant and mice of *Tor1a* knockout or *Tor1a*^*ΔE/ΔE*^ ^10,14,37^. This discrepancy may be caused by species differences and functional redundancy between different human torsin genes in NE morphology maintenance ^11,51^. Only one torsin gene exists in *Drosophila* but there are 4 torsin genes in humans. Additionally, all human patients are heterozygous with one remaining WT copy of *TOR1A* gene that may also diminish such severe abnormalities as nuclear blebs. As such, the impaired mRNA export in DYT1 MNs might be mainly due to the defective NPC-mediated transport pathway, rather than the NE-budding mechanism ^32^.

### The Thickened Nuclear Lamina and Impaired NCT

The nuclear envelope and embedded NPCs play critical roles in regulating nucleocytoplasmic transport of both transcripts and proteins ^52–55^. The impaired mRNA nuclear export and protein NCT in DYT1 MNs indicate some fundamental factors are dysregulated. Our study indicates that NE abnormality including the thickened nuclear lamina may contribute to the impaired NCT in DYT1 MNs (Fig. 8I). Mislocalized and dysfunctional NPCs lacking the NUP358 subunit appear in *Tor1a*-null mouse neurons ^56^. Nonetheless, we failed to detect significant changes on NPCs in human DYT1 neurons. The protein levels of NUP98 and NUP358 are not significantly altered; and the signal densities of nuclear pores probed by antibodies for NUP358, NUP98 and other NPC proteins (MAB414) are also not significantly different from controls. The NPC size in human DYT1 MNs is also similar to that in control cells when examined under TEM. These results indicate that the overall NPC number and composition may not be significantly altered in human DYT1 MNs. However, NPC function could be disrupted by the thickened nuclear lamina and abnormal NE morphology (Fig. 8I).

Defective NCT is emerged as a common molecular underpinning for many neurological diseases ^57–60^. Impaired NPCs and dysregulated NCT signaling pathways are the focus of many prior investigations. Our findings provide a new angle to understand impaired NCT. NE abnormalities including the thickened nuclear lamina and disrupted morphology may be fundamental to many diseases with defective transports.

## Materials and methods

### Cell lines and culture condition

All cell lines used in this study are listed in Supplemental Table S1. HEK 293T cells and SH-SY5Y cells were purchased from ATCC. DYT1 fibroblast and healthy controls were obtained from the *Coriell Institute for Medical Research*. Human age-matched WT iPSC was obtained from Coriell and H9 ESC line was purchased from *WiCell Research Institute*. Human iPSCs were maintained in complete mTeSR1 medium (STEMCELL Technologies) on Matrigel (Corning) coated dishes at 37°C and 5% CO_2_, and the medium was daily replaced. The medium recipes were provided in supplemental information.

### Plasmid construction and virus production

A third-generation lentiviral vector (*pCSC-SP-PW-IRES-GFP*) was used to express *NEUROG2-IRES-GFP-T2A-Sox11*, *NEUROG2-IRES-Sox11*, and *ISL1-T2A-LHX3*. cDNAs for *TOR1A* and *TOR1AΔE* were provided by Dr. Gonzalo E. Torres ^17^ and were individually subcloned into *pCSC-SP-PW-IRES-GFP* vector. Another third-generation lentiviral vector (*LV-CAG-mCherry-miRE-Luc*) was used to express *TOR1A*-shRNAs and *LMNB1*-shRNAs as described in previously report ^61^. The target sequences of shRNAs are listed in supplemental Table S3. The dual reporter 2Gi2R was generously provided by Dr. Fred H. Gage ^35^. For retrovirus package, four moloney-based retroviral vectors (pMXs) carrying human complementary DNAs of *OCT4*, *SOX2*, *KLF4* and *c-MYC* were obtained from Addgene ^62^. Each of these plasmids was co-transfected with two packaging vectors (*pCMV-Gag-Pol* and *pCMV-VSVG*) into HEK293T cells. Replication-incompetent lentiviruses were produced and viral supernatants were collected at 48 hrs and 72 hrs post-transfection ^63^. The viral supernatants were filtered through 0.45 μm syringe filters and stored at 4 °C prior to cell transduction.

### Direct conversion of adult fibroblasts into MNs

Lentiviral delivery of 4 factors (*NEUROG2*, *Sox11*, *ISL1* and *LHX3*) was used as previously described ^23,24^. In brief, fibroblasts were plated at a density of 1×10^4^ cells/cm^2^ onto Matrigel-coated dishes. Cells were transduced the next day with lentiviral supernatants supplemented with 6 mg/ml polybrene. Fibroblast medium was refreshed after overnight incubation. One day later, the culture was replaced with neuronal induction medium and half-changed every the other day until 14 dpi (Fig 1A). A replating procedure ^64^ was used to purify induced neurons. The purified cells were cultured in neuronal maturation medium onto Matrigel-coated coverslips with or without the presentence of astrocytes depending on desired experiments. The medium was half changed twice a week until analysis.

### Immunostaining and confocal microscopy

Cultured cells at indicated time points were fixed with 4% paraformaldehyde (PFA) in PBS for 15 min at room temperature and then incubated in blocking buffer (PBS containing 0.2% Triton X-100 and 3% BSA) for 1 hour for permeabilization and blocking. Cells were then incubated with primary antibodies in blocking buffer at 4°C overnight and then followed by washing and incubation with corresponding fluorophore-conjugated secondary antibodies. All antibodies used in this study are listed in Supplemental Table S4. The nuclei were stained with Hoechst 33342 (HST, ThermoFisher Scientific).

Confocal images were obtained with a Nikon A1R or Zeiss LSM700 (63×/1.4 NA) Confocal Microscopes. Neurite length was measured by ImageJ as previously described ^27–29^. For nuclear morphology assay, we zoomed in individual nucleus using ImageJ software and defined the abnormal nucleus based on two criteria: 1) at least one obvious sharp angle or a deep invagination, and 2) at least half area of the nucleus losing the smooth outline. Signals of nuclei (HST) or nuclear lamins were used to distinguish the nucleus and cytoplasm.

### Fluorescent in situ hybridization (FISH)

*In situ* hybridization was performed as described previously ^25^. Briefly, cultured cells were washed once with PBS, and fixed with 4% PFA for 30 min at room temperature. Cells were further permeabilized with 0.2% TritonX-100 for 10 min and washed with PBS twice afterwards. Samples were equilibrated with hybridization buffer composed of 2XSSC, 10%dextran sulfate, 10 mM ribonucleoside-vanadyl complex (RVC; New England Biolabs) and 20% formamide. A mixture of d*igoxigenin* (*DIG*)-labeled oligo-dT or dA probes (0.2ng/μL) (Table S3) and yeast tRNA (0.2μg/μL) were heated to 95°C for 5min and chilled on ice immediately. Probes were then combined with equal volumes of 2X hybridization buffer and added to samples for incubation at 37°C overnight. Anti-*DIG* antibody (Sigma) and fluorophore-conjugated corresponding secondary antibodies (Thermo Fisher Scientific) were used to detect oligo-dT signals. Anti-MAP2 (Abcam) antibody and Hoechst 33342 dye were used to determine the neuron soma and nuclei, respectively. Confocal images were zoomed in using ImageJ software to measure the fluorescent signal density. The “mean” values of the nucleus and the cytoplasm in each neuron were used to calculate the ratio of dT signal in nucleus to cytoplasm ^25^. We measured as large an area as possible for each neuron and employed an unbiased approach for data collection. The person who analyzed the images was completely blinded to the sample information.

### Western blot analysis

Induced iPSC-MNs were lysed in lysis buffer composed of 50 mM Tris-HCl buffer (pH 8.0), 150 mM NaCl, 1% NP40, 1% Triton X-100, 0.1% SDS, 0.5% sodium deoxycholate, and protease inhibitor cocktail (Roche). Equal amounts of these lysates (20 mg per lane) were used for SDS-PAGE and western blot analysis as previously described ^29^. Primary antibodies and dilutions are listed in supplemental Table S4.

### Quantitative real-time PCR analysis

Total RNA was extracted from cultured cells using TRIzol (Life Technologies), and genomic contamination was removed using TURBO DNase (Life Technologies). cDNA synthesis reactions were performed using 0.5 μg of RNA from each sample with the SuperScriptIII First-Strand kit (Life Technologies) and random hexamer primers. Real-time PCR was performed in triplicate using primers, SYBR GreenER SuperMix (Invitrogen), and the ABI 7900HT thermocycler (Applied Biosystems). Primers of RT-PCR assay are listed in supplemental Table S3. Target mRNA levels were normalized to the reference gene *HPRT* by 2^−ΔΔCt^ method as described previously ^27–29^. PCR primer sequences are available upon request.

### Transmission electron microscopy (TEM)

Reprogrammed neurons were cultured onto Matrigel-coated glass coverslips and fixed at desired time points. The fixative solution was composed of 4% paraformaldehyde, 1% glutaraldehyde, 1.5 μm MgCl_2_ and 5 μm CaCl_2_ in 0.1 M cacodylate buffer (pH7.4) and the fixation was performed at 4°C overnight. Samples were washed with cacodylate buffer twice and treated with 1% osmium tetraoxide (OsO_4_, EMS) with 0.8% K3Fe(CN6) in cacodylate buffer at room temperature for 1.5 hours, and then prestained with 2% uranyl acetate in water for 2 hours at room temperature in the dark. The following steps of dehydration, infiltration, and embedding were the same as previously described ^34^. The resin blocks were polymerized at 60°C overnight and sectioned with a diamond knife (Diatome) on a Leica Ultracut UCT 6 ultramicrotome (Leica Microsystems). Sections were post stained with 2% uranyl acetate in water and lead citrate. Images were acquired on a JEOL JEM 1200EX transmission electron microscope equipped with a tungsten source at 120kV using a Morada SIS camera (Olympus). The electron-dense areas immediately beneath the nuclear membrane were measured by ImageJ (NIH) and the ratio of electron-dense area to the nuclear perimeter was used to evaluate the thickness of nuclear lamina. The number and the size of nuclear pore complexes were quantified by ImageJ (NIH) as previously described.

### Statistical analysis

Unpaired two-tailed Student’s *t* tests were used to compare one experimental sample with its control. One-way analysis of variance was used when comparing multiple samples, with either Tukey (when comparing samples with each other) or Dunnett (when comparing samples to a control) post hoc tests. Results are expressed as mean ± SD of at least three biological replicates, and *P* < 0.05 was treated as significant.

Methods of DYT1 iPSC generation, neural progenitor generation, neural differentiation, and electrophysiology were described in supplemental information.

## Supporting information

Supplemental Materials including supplemental text, figures and tables

## Author contributions

Conceptualization, B.D., Y.T. and C.L.Z.; Methodology, B.D., Y.T. and M.L.L.; Investigation, B.D., Y.T., S.M., T. Z. and M.A.; Writing – Original Draft, B.D.; Writing – Review & Editing, B.D., Y.T. and C.L.Z.; Funding Acquisition, B.D., Y.T. and C.L.Z.; Supervision, B.D. and C.L.Z.

## Acknowledgments

We thank members of the Zhang laboratory (Xiaoling-Zhong, Wenjiao Tai, Leilei Wang, Yunjia Zhang and Yuhua Zou) and the Ding laboratory (Masood Sepehrimanesh, Haochen Cui and Jacob A Stagray) for technical help and Dr. William T. Dauer (UT Southwestern) for discussion. We also appreciate the support from the UT Southwestern Electron Microscopy Core and the Microscopy Center at the University of Louisiana at Lafayette. This work was supported by The Welch Foundation (I-1724), The Mobility Foundation, The Decherd Foundation, The Pape Adams Foundation, the Kent Waldrep Foundation Center for Basic Research on Nerve Growth and Regeneration, National Institutes of Health (NIH) grants (NS092616, NS099073, and NS088095 to C.L.Z.; R21NS112910 to B.D.), and Friends of the Alzheimer’s Disease Center (ADC) and NIH ADC grant (NIH/NIA P30-12300-21 to B.D.) and the Department of Defense Peer Reviewed Medical Research Program (PRMRP) grant (W81XWH2010186 to B.D.), and National Natural Sciences Foundation of China (No. 81801200 to Y.T.) and Hunan Provincial Natural Science Foundation of China (No. 2019JJ40476 to Y.T.).

## Conflict of interest statement

The authors declared that no conflict of interest exists.

## Notes

### Competing Interest Statement

The authors have declared no competing interest.

