## Supplemental Materials including supplemental text, figures and tables for "Disease modeling with human neurons reveals LMNB1 dysregulation underlying DYT1 dystonia"

**This PDF file includes:**
Supplementary information text
Figures S1-S8
Table S1-S4
References

**Supplementary information text**

**Materials and Methods**

***Cell lines and culture condition***

All cell lines used in this study are listed in Supplemental Table S1. HEK 293T cells and SH-SY5Y cells were purchased from ATCC. DYT1 fibroblast and healthy controls were obtained from the Coriell Institute for Medical Research. Human age-matched WT iPSC was obtained from Coriell and H9 ESC line was purchased from WiCell Research Institute. Human iPSCs were maintained in complete mTeSR1 medium (STEMCELL Technologies, Vancouver, BC, Canada) on Matrigel (Corning) coated dishes at 37°C and 5% $CO_2$, and the medium was daily replaced.

The medium recipes were:

(1) Fibroblast medium: DMEM supplemented with 15% fetal bovine serum (FBS, Corning, NY, USA) and 1% penicillin/streptomycin.

(2) Neuronal induction medium: DMEM:F12:neurobasal (2:2:1), 0.8% N2 (Invitrogen), 0.8% B27 (Invitrogen), 1% penicillin/streptomycin, and supplemented with 10 mM FSK (Sigma-Aldrich), 1 mM dorsomorphin (DM, Millipore, MA, United States) and 10 ng/ml bFGF (PeproTech).

(3) Neuronal maturation medium: DMEM:F12:neurobasal (2:2:1), 0.8% N2 (Invitrogen), 0.8% B27 (Invitrogen), 1% penicillin/streptomycin, and supplemented with 5 mM FSK and 10 ng/ml each of BDNF, GDNF, and NT3 (PeproTech).

(4) ESC medium: DMEM/F12 medium with 20% KnockOut Serum Replacement (KOSR, Thermo Fisher Scientific, MA, USA), 1% GlutaMax, 1% non-essential amino acid (NEAA), 50 μM β-mercaptoethanol (β-ME), 1% P/S and 10 ng/ml basic fibroblast growth factor (bFGF, PeproTech, NJ, USA).

(5) KOSR medium: DMEM/F12 medium with 20% KOSR, 1% GlutaMax, 1% NEAA, 50 μM β-ME and 1% P/S.

(6) Neurosphere medium (NSP medium): DMEM/F12 medium containing 1% N2, 1% GlutaMax, 1% NEAA, 50 μM β-ME, 1% P/S, 8 μg/ml Heparin, 20 ng/ml bFGF and 20 ng/ml epidermal growth factor (EGF, PeproTech).

(7) Neural progenitor cell medium: DMEM/F12 and neurobasal medium (1:1) containing 0.5% N2 (Invitrogen, CA, USA), 1% B27 (Invitrogen), 1% GlutaMax, 1% NEAA, 50 μM β-ME, 1% P/S, 10 ng/ml EGF and 10 ng/ml bFGF.

(8) SH-SY5Y induction medium: DMEM supplemented with 3% FBS, 10 μM all-trans-Retinoic acid (Sigma) and 1% penicillin/streptomycin.

***Plasmid construction and virus production***

A third-generation lentiviral vector (*pCSC-SP-PW-IRES-GFP*) was used to express *NEUROG2-IRES-GFP-T2A-Sox11*, *NEUROG2-IRES-Sox11*, and *ISL1-T2A-LHX3*. cDNAs for *TOR1A* and *TOR1AΔE* were provided by Dr. Gonzalo E. Torres [1] and were individually subcloned into *pCSC-SP-PW-IRES-GFP* vector. Another third-generation lentiviral vector (*LV-CAG-mCherry-miRE-Luc*) was used to express *TOR1A*-shRNAs and *LMNB1*-shRNAs as described in previously report [2] The dual reporter 2Gi2R was generously provided by Dr. Fred H. Gage [3]. For retrovirus package, four moloney-based retroviral vectors (pMXs) carrying human complementary DNAs of *OCT4*, *SOX2*, *KLF4* and *c-MYC* were obtained from Addgene (Takahashi et al., 2007). Each of these plasmids was co-transfected with two packaging vectors (*pCMV-Gag-Pol* and *pCMV-VSVG*) into HEK293T cells (ATCC). Replication-incompetent lentiviruses were produced and viral supernatants were collected at 48 hrs and 72 hrs post-transfection [4]. The viral supernatants were filtered through 0.45 μm syringe filters and stored at 4 °C prior to cell transduction.

***Direct conversion of adult fibroblasts into MNs***

Lentiviral delivery of 4 factors (*NEUROG2*, *Sox11*, *ISL1* and *LHX3*) was used as previously described [5,6]. In brief, fibroblasts were plated at a density of $1x10^4$ cells/$cm^2$ onto Matrigel-coated dishes. Cells were transduced the next day with lentiviral supernatants supplemented with 6 mg/ml polybrene. Fibroblast medium was refreshed after overnight incubation. One day later, the culture was replaced with neuronal induction medium and half-changed every the other day until 14 dpi (Fig 1A). A replating procedure [7] was used to purify induced neurons. The purified cells were cultured in neuronal maturation medium onto Matrigel-coated coverslips with or without the presentence of astrocytes depending on desired experiments. The medium was half changed twice a week until analysis.

***Immunostaining and confocal microscopy***

Cultured cells at indicated time points were fixed with 4% paraformaldehyde (PFA) in PBS for 15 min at room temperature and then incubated in blocking buffer (PBS containing 0.2% Triton X-100 and 3% BSA) for 1 hour for permeabilization and blocking. Cells were then incubated with primary antibodies in blocking buffer at 4°C overnight and then followed by washing and incubation with fluorophore-conjugated corresponding secondary antibodies. All antibodies used in this study are listed in Supplemental Table S3. The nuclei were stained with Hoechst 33342 (HST, ThermoFisher Scientific).

Confocal images were obtained with a Nikon A1R or Zeiss-LSM700 Confocal Microscopes. Neurite length was measured by ImageJ as previously described [8-10]. For nuclear morphology assay, we zoomed in individual nucleus using ImageJ software and defined the abnormal nucleus based on two criteria: 1) at least one obvious sharp angle or a deep invagination, and 2) at least half area of the nucleus losing the smooth outline. Signals of nuclei (HST) or nuclear lamins were used to distinguish the nucleus and cytoplasm.

***DYT1 iPSC generation***

DYT1 fibroblasts were plated at $8x10^5$ cells/dish onto gelatin-coated 10-cm dishes, and were inducted with the cocktail of four retroviruses (1:1:1:1) expressing *OCT4*, *SOX2*, *KLF4* and *c-MYC*. Fibroblasts were received a second round of virus infection two days later, and was marked as day 0. Fibroblasts were replated at day 5 with fibroblast medium at $2x10^5$ cells per well in Matrigel-coated 6-well plates. The medium was replaced with human ESC medium at day 6, and then changed as mTeSR1 medium in the following few days. Cells were treated with 0.5 mM VPA (Sigma) at day 5 and lasted for 10 days. iPSC colonies were picked around 3 weeks to 1 month post-induction based on ESC-like colony morphology. The picked colonies were then expanded and maintained on Matrigel-coated plates with mTeSR1 medium until characterization.

***Embryoid bodies (EBs)***

hPSCs were dissociated by Versene (Thermo Fisher Scientific) treatment and transferred to low attachment 10-cm petri dishes (Grainger, IL, USA) in KOSR medium supplemented with 10 μM ROCK inhibitor Y-27632 (STEMCELL Technologies). The medium was changed every other day. Seven days after suspension culture, EBs were digested with 0.25% Trypsin and cultured on gelatin-coated plates with KOSR medium for another seven days. RNAs were eventually isolated and used for the qRT-PCR analysis of three germ layers.

***Neural progenitor cells generation and neural differentiation***
Neural progenitor cells were generated as previously reported with minor modifications [11]. Briefly, hPSCs were cultured in mTeSR1 medium with 10 µM all-trans-retinoic acid (RA, Sigma) and 0.5 mM VPA on Matrigel-coated 6-cm plates for seven days. hPSCs were then digested with Versene and gently pipetted into small clumps supplemented with 10 µM Y-27632. Cell clumps were aggregated in KOSR medium for four days, followed by culturing in NSP medium for another one week. Neurospheres were then formed and dissociated into single cells by accutase (Innovative Cell Technologies, CA, USA), and finally maintained in neural progenitor cell medium.

For MN differentiation, neural progenitor cells were plated onto Matrigel-coated plates at a density of $3x10^4$ cells/$cm^2$ and inducted with a cocktail of two lentiviruses (1:1) expressing *NEUROG2-IRES-GFP-T2A-Sox11* and *ISL1-T2A-LHX3* respectively. The next day, the medium was replaced with neuronal maturation medium. Neurons were then dissociated by Accutase at day 6 and replated onto astrocyte-coated coverslips for long-term culture.

***Alkaline phosphatase (AP) staining and karyotyping***
AP staining was performed with the Vector Blue Substrate kit (SK-5300) from Vector Laboratories. Briefly, iPSCs were fixed with 4% paraformaldehyde (PFA) supplemented with 0.1% Triton X-100 for 10 min. After rinses with PBS and $H_2O$, cells were then incubated with freshly prepared AP working solution in the dark for 40 min. Karyotype analysis of iPSCs was performed at WiCell Research Institute.

***Bisulfite sequencing and T-A sequencing***
Genomic DNA from fibroblasts and hPSCs were extracted with lysis buffer containing 0.5 mg/ml proteinase K. Bisulfite conversion of genomic DNA was carried out using the EZ DNA Methylation-Direct Kit (Zymo Research) according to the manufacturer's instructions. A genomic fragment of the *OCT4* promoter was amplified using EpiMark Taq DNA polymerase (New England Biolabs). Primer sequences were previously published [12]. PCR products were then purified by gel extraction using Zymoclean Gel DNA Recovery Kit (Zymo Research), and subsequently cloned into the pMD20-T vector (Takara, Shiga, Japan). Single clones were picked from each sample (n=7-11/sample) and sequenced by the M13Rev universal primer.

To confirm the mutations of *TOR1A* in generated iPSCs, genomic DNA were similarly isolated from WT and DYT1 iPSC, amplified by PCR and cloned into the *pMD20-T* vector. Single clones were picked from each sample (n=4/sample) and sequenced by the *M13Rev* universal primer.

***Teratoma generation***
Briefly, around 5 million iPSCs were injected into testis capsules of immunocompromised *NOD/SCID* mice (aged 2-3 months). Tumor samples were collected in 8-12 weeks, fixed in 10% neutral buffered formalin (NBF), and processed for paraffin embedding and hematoxylin and eosin (H&E) staining. Experimental protocols were approved by the Institutional Animal Care and Use Committee at University of Texas Southwestern.

***Electrophysiology***

Reprogrammed MNs were cultured on astrocyte-coated glass coverslips for 35 days and used for analysis. Whole-cell patch-clamp recordings were performed under visual guidance using GFP-fluorescence to identify GFP + cells when cells were maintained at 30 °C in a submersion chamber with Tyrode solution containing 150 mM NaCl, 4 mM KCl, 2 mM MgCl2, 3 mM CaCl2, 10 mM glucose, and 10 mM HEPES at pH 7.4 (adjusted with KOH) and 300 mOsm. The recording pipettes (approximately 6–9 MΩ) were filled with an intracellular solution containing 0.2 mM EGTA, 130 mM K-gluconate, 6 mM KCl, 3 mM NaCl, 10 mM HEPES, 4 mM ATP-Mg, 0.4 mM GTP-Na, 14 mM phosphocreatine-di(Tris) at pH 7.2 and 285 mOsm. Series and input resistance were measured in voltage-clamp with a 400 ms, 10 mV step from a -60 mV holding potential (filtered at 30 kHz and sampled at 50 kHz). Cells were used for analysis only if the series resistance was less than 30 MΩ and was stable throughout the experiment. The input resistance ranged from 0.2 to 2 GΩ. Currents were filtered at 3 kHz, acquired and digitized at 10 kHz using Clampex10.3 software (Molecular Devices). Action potentials were recorded in current-clamp and elicited by a series of current injections ranging from -20 to 200 pA at 20 pA increments and 800 ms in duration. All current-clamp recordings were made at resting membrane potential or without any current injection. Data analysis was performed with Clampfit10.3 software (Molecular Devices). The action potential (AP) trace immediately above threshold was used to determine the delay of 1$^{st}$ spike as the length of time and the start of current steps to the peak of AP. The same AP trace was used to measure AP threshold as the corresponding voltage when there was the sharpest change of the trace slope. The above indicated AP trace was also measured to determine maximum velocity of the rise and decay. AP frequency was obtained by dividing the maximum number of spikes during the current steps protocol with the step time duration (800ms).

***Statistical analysis***

Data were presented as means ± SEM and analyzed by GraphPad Prism 6.0 (GraphPad Software, Inc., La Jolla, CA, USA). Statistical analysis was conducted by one-way analysis of variance and statistical significance was set at p-value < 0.05.

**Supplementary Figures and Figure Legends**

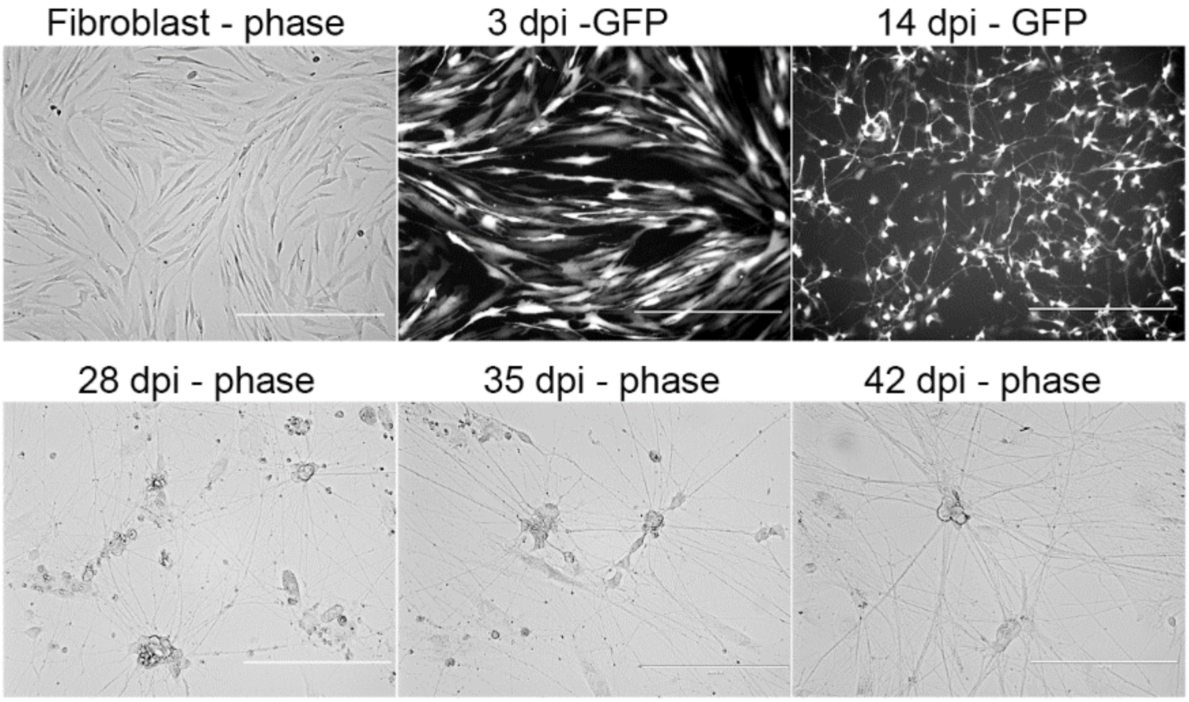


**Figure S1 (related to Figure 1). Direct conversion of adult fibroblasts into MNs.** Micrographs of fibroblasts and diMNs at the indicated time points. Scale bar: 200 µm.

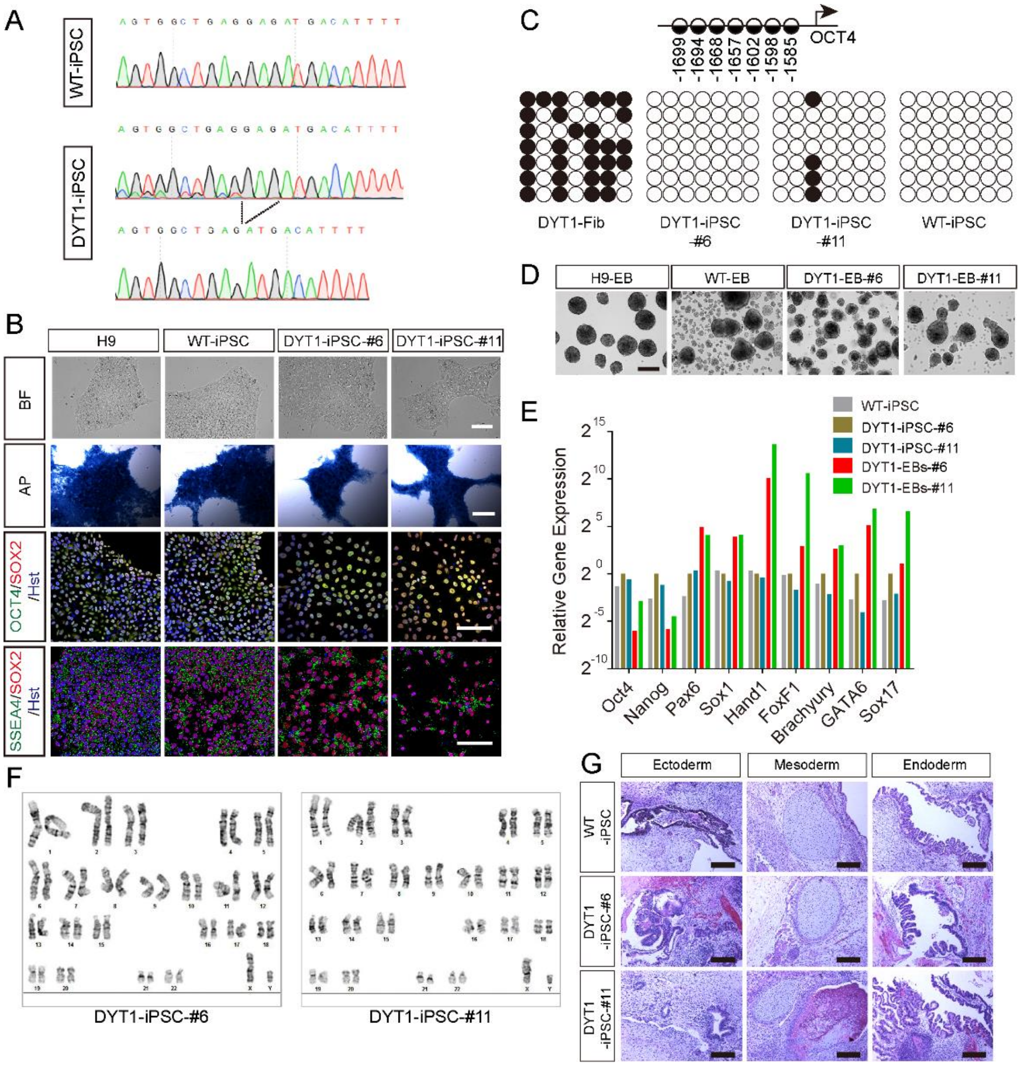


**Figure S2 (related to Figure 3). Generation and characterization of DYT1 iPSCs.**
**(A)** Sequence confirmation of *TOR1A* mutation in DYT1 iPSCs.
**(B)** Pluripotency of iPSCs. Bright-field images (Scale bar, 100 μm), alkaline phosphatase (AP) staining (Scale bar, 250 μm) and immunostaining of pluripotency markers OCT4, SOX2 and SSEA4 in iPSCs (Scale bars, 100 μm).
**(C)** Methylation state of the promoter region of OCT4. Positions of the CpG dinucleotides relative to the OCT4 transcription start site is provided. Open and closed circles indicate unmethylated and methylated CpGs, respectively.
**(D)** Generation of embryonic bodies (EBs) from iPSCs in suspension culture at day 7. Scale bars, 200 μm.
**(E)** qRT-PCR analysis of endogenous ESC markers in iPSCs and three germ layer markers in EBs at day 14.
**(F)** Karyotypes of DYT1-iPSCs.
**(G)** H&E staining of teratoma sections showed differentiation of iPSCs to various tissues of three germ layers, such as neural epithelium, cartilage, muscle and glandular structures. Scale bars, 250 μm.

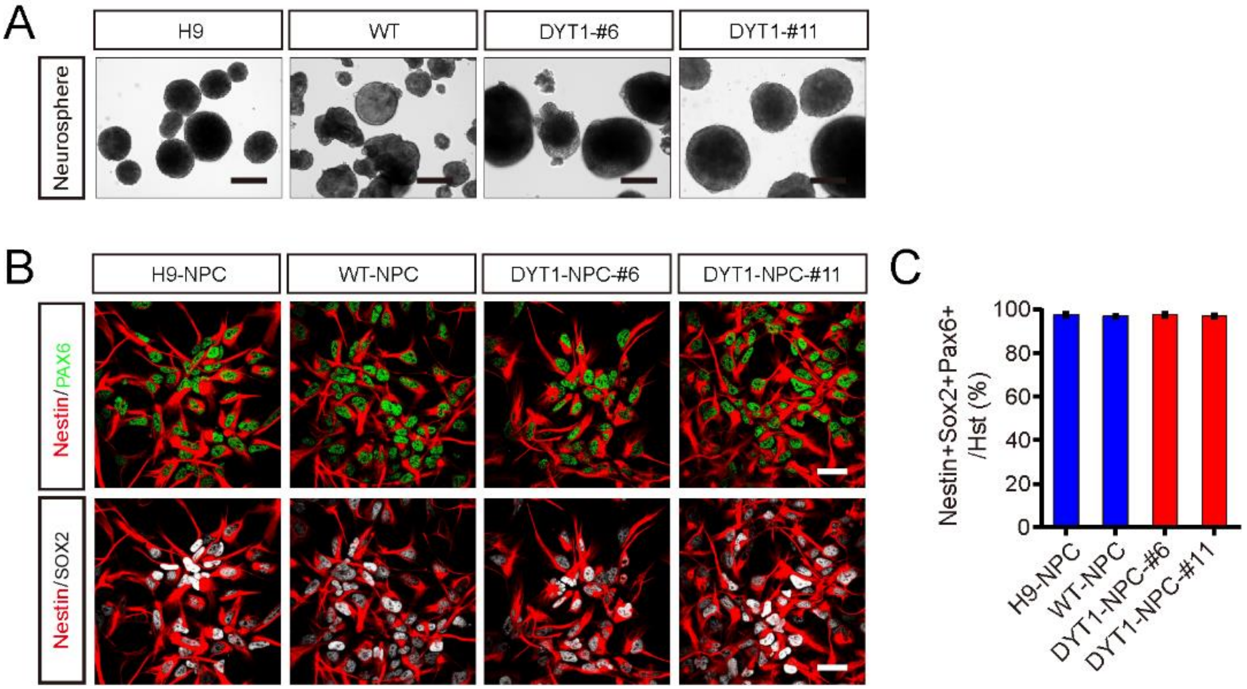


**Figure S3 (related to Figure 3). Generation of neural progenitor cells.**
**(A)** Neurosphere formation at day 18. Scale bar, 200 µm.
**(B, C)** Immunostaining of neural progenitor cell markers. Scale bar, 50 µm. The ratios of nestin, SOX2 and PAX6 triple positive cells were calculated respectively as the purity of generated neural progenitor cells (n = 204 for H9, n = 233 for WT, n = 237 for DYT1-#6, and n = 238 for DYT1-#11 from triplicates).

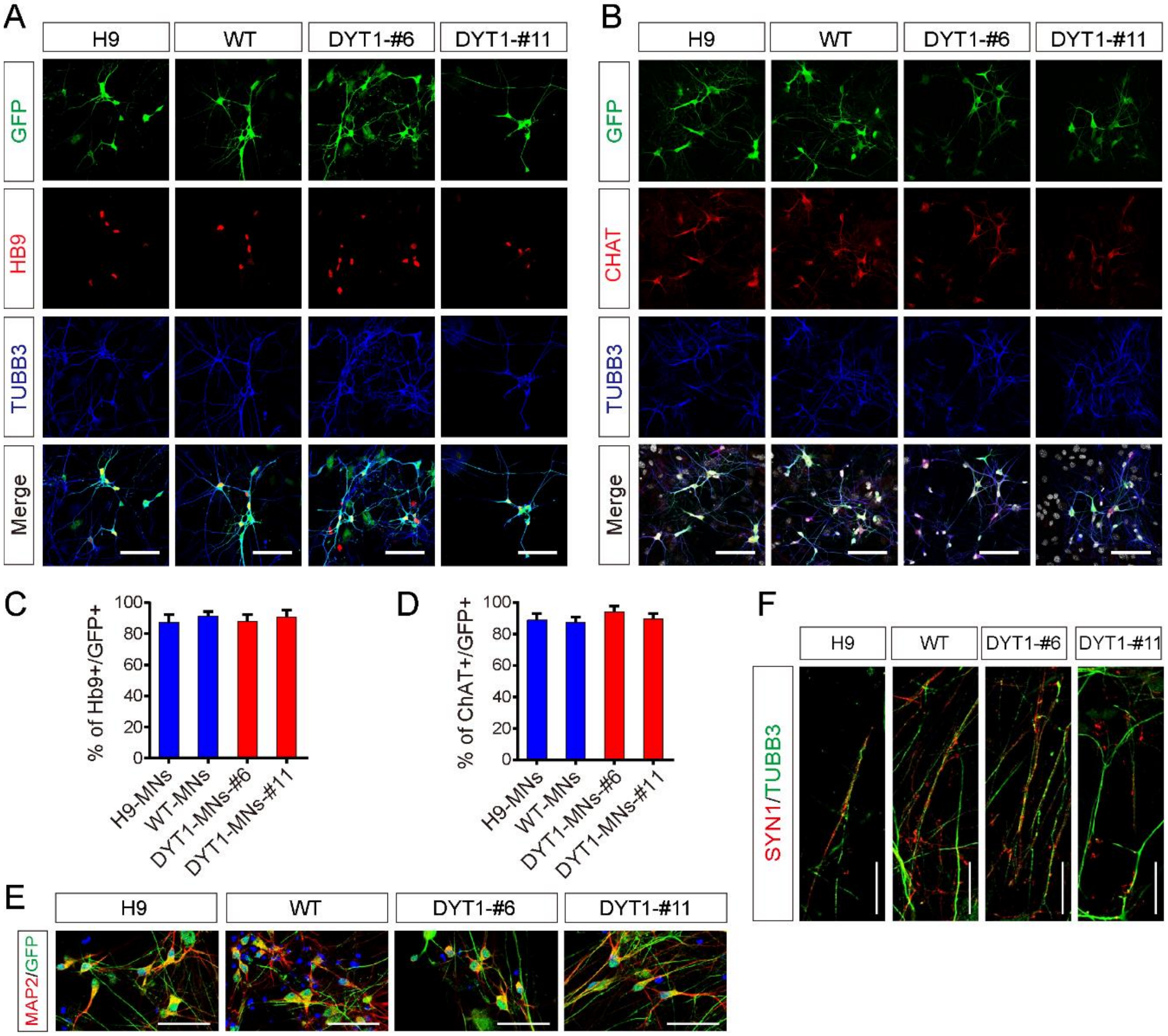


**Figure S4 (related to Figure 3). Differentiation of MNs from iPSCs.**

**(A, B)** Immunostaining of MN markers, HB9 and CHAT at 3 wpi. Scale bars, 100 µm.

**(C, D)** The ratios of $HB9^{+}/GFP^{+}$ neurons (n = 132 for H9, n = 216 for WT, n = 152 for DYT1-#6, and n = 120 for DYT1-#11) and $CHAT^{+}/GFP^{+}$ neurons (n = 160 for H9, n = 252 for WT, n = 112 for DYT1 -#6, and n = 168 for DYT1-#11) were calculated respectively.

**(E)** Immunostaining of a mature neuron marker MAP2 at 3 wpi. Scale bars, 50 µm.

**(F)** Immunostaining of a pre-synaptic marker Synapsin 1 (SYN1) at 3 wpi. Scale bars, 25 µm.

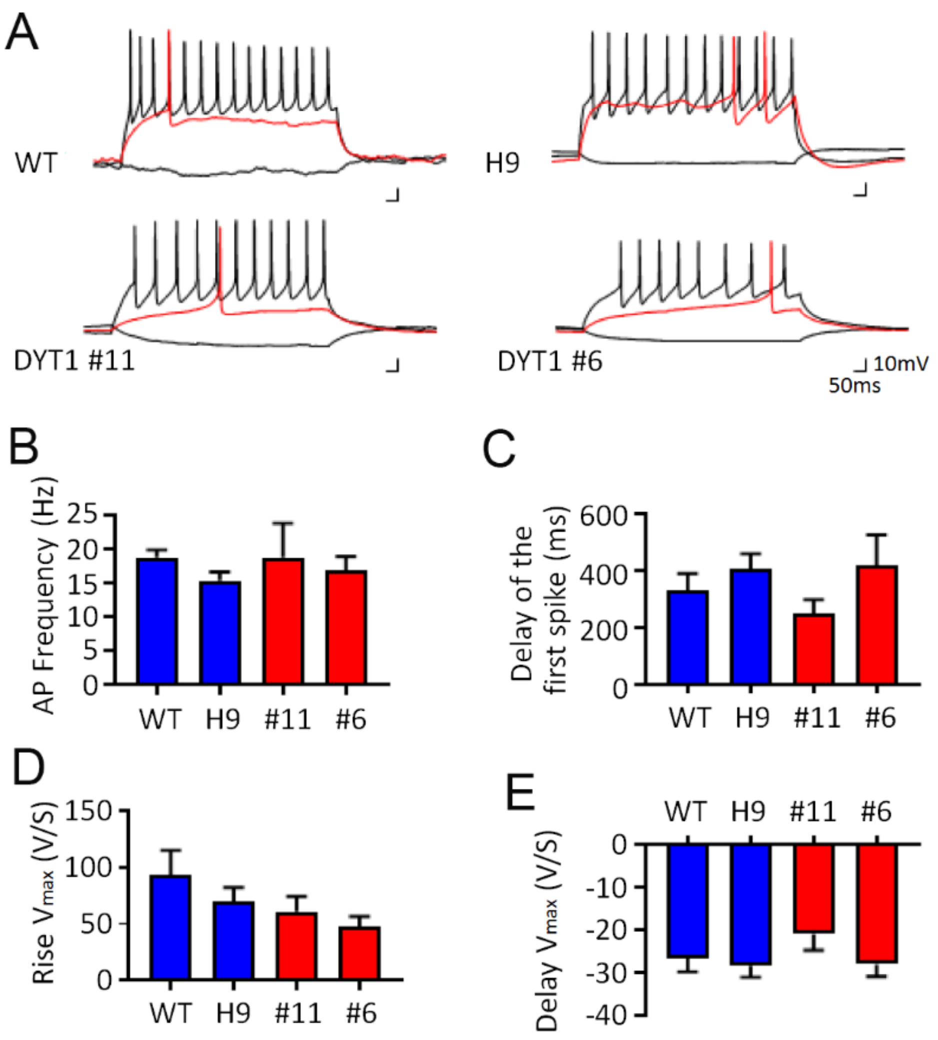


**Figure S5 (related to Figure 3). Electrophysiological analysis of iPSC MNs.**
**(A)** Repetitive action potential (AP) waveforms recorded under current-clamp mode for the control (WT and H9) and DYT1 iPSC-MNs at 5 wpi. The precondition sweep and the sweep immediately above threshold (in red) are also shown.
**(B-E)** Quantifications of AP frequency, delay of the first spike, rise and delay $V_{max}$ in indicated cells, respectively (n = 30 for each line from 4 biological replicates).

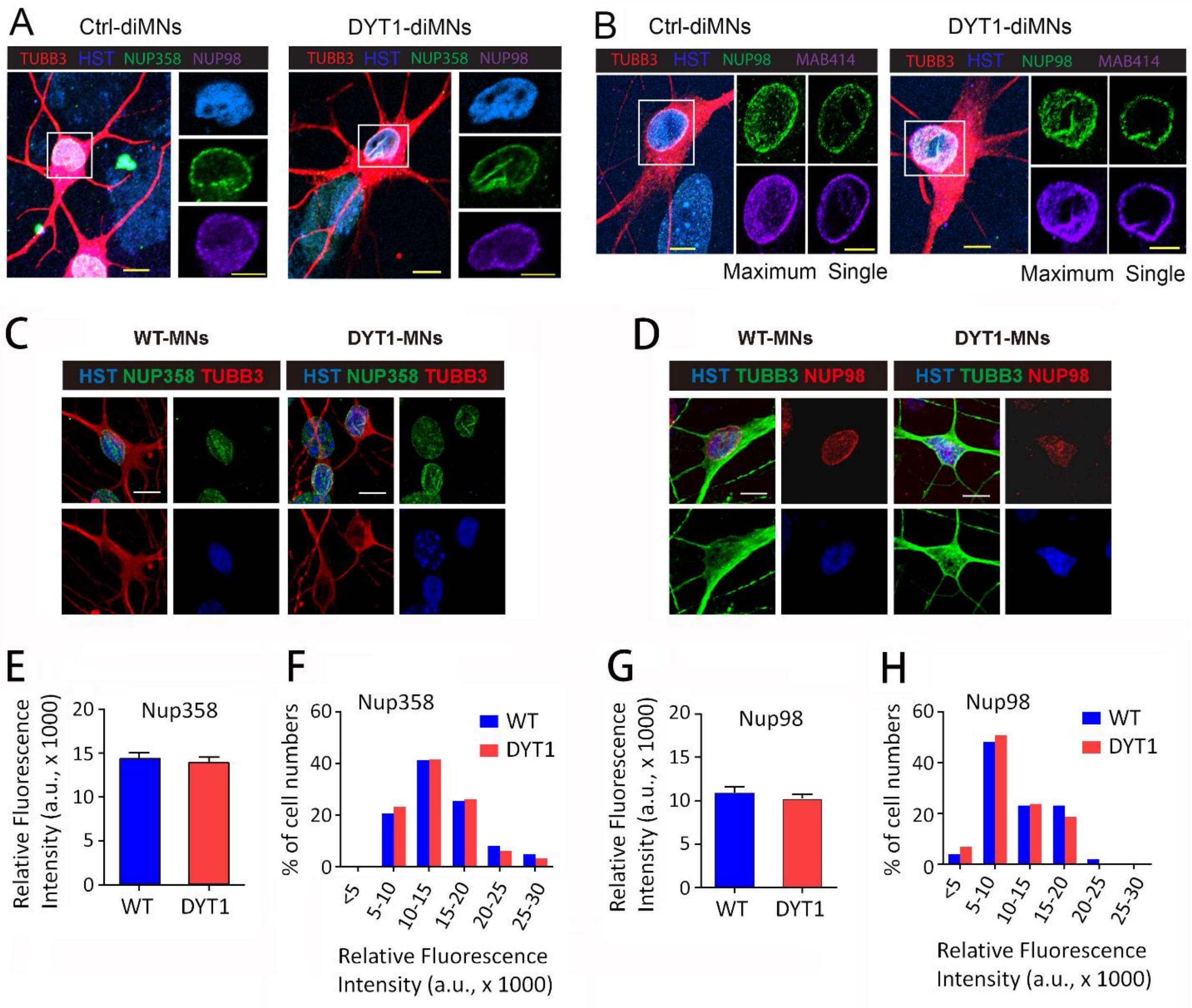


**Figure S6 (related to Figures 4 and 7). Immunocytochemistry of nuclear pore complex subunits in DYT1-MNs.**

**(A)** NUP358 and NUP98 expression in control and DYT1-diMN at 6 wpi. Scale bars: 20 μm.

**(B)** NUP98 and MAB414 examined nuclear pore complex in control and DYT1-diMN at 6 wpi. Scale bars: 20 μm.

**(C)** NUP358 expression in WT and DYT1 iPSC-MN at 3 wpi. Scale bars: 10 μm.

**(D)** NUP98 expression in WT and DYT1 iPSC-MN at 3 wpi. Scale bars: 10 μm.

(**E-F**) Relative fluorescence intensity of NUP358 and its intensity distributions in WT and DYT1 iPSC-MN at 3 wpi. n (neurons) >100 each samples from triplicates.

(**G-H**) Relative fluorescence intensity of NUP98 and its intensity distributions in WT and DYT1 iPSC-MN at 3 wpi. n (neurons) >100 each samples from triplicates.

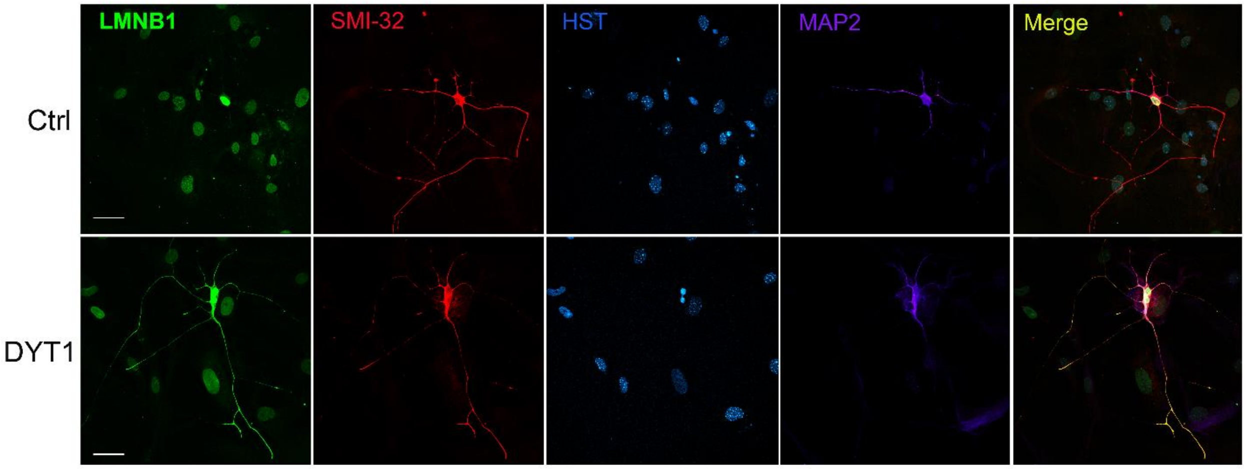


**Figure S7 (related to Figure 5). LMNB1 mislocalization in both axons and dendrites of DYT1 diMNs.**
Confocal images of DYT1 diMNs at 6 wpi. Scale bar: 50 µm.

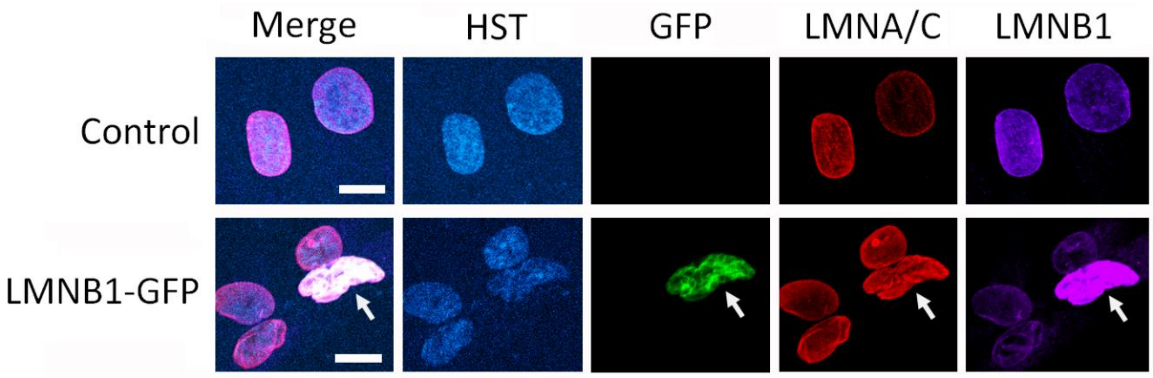


**Figure S8 (related to Figures 7 and 8). Overexpression of LMNB1 disrupts nuclear envelope morphology.** Confocal micrographs of fibroblasts with or without overexpression of LMNB1 (fused with GFP). Arrow indicates a cell with LMNB1 overexpression. Scale bars, 10 µm.

**Table S1 List of cell lines used in this study and their Research Resource Identifiers.**

| | Identifiers (RRIDs) | Sex | Age at Sampling (YR) | Tissue Type | Race | Onset Age (YR) | Gene | Mutation |
|---|---|---|---|---|---|---|---|---|
| **Fibroblast** | | | | | | | | |
| DYT1-1 | Coriell Cat# GM03211, RRID:CVCL_1U24 | M | 30 | Skin | Caucasian | 7 | TOR1A | 907_909delGAG |
| DYT1-2 | NDS00305 | M | 22 | Skin | Caucasian | 12 | TOR1A | 907_909delGAG |
| DYT1-3 | Coriell Cat# GM02304, RRID:CVCL_1U21 | F | 17 | Skin | Caucasian | 7 | TOR1A | 907_909delGAG |
| DYT1-4 | NDS00301 | F | 53 | Skin | Caucasian | 9 | TOR1A | 907_909delGAG |
| Ctrl-1 | Coriell Cat# GM00024, RRID:CVCL_7269 | M | 31 | Skin | Caucasian | | | |
| Ctrl-2 | Coriell Cat# GM03652, RRID:CVCL_7397 | M | 24 | Skin | Caucasian | | | |
| Ctrl-3 | Coriell Cat# GM04506, RRID:CVCL_7413 | F | 20 | Skin | Caucasian | | | |
| Ctrl-4 | Coriell Cat# AG07473, RRID:CVCL_2C33 | F | 50 | Skin | Caucasian | | | |
| **Induced pluripotent stem cells (control)** | | Coriell Cat# GM23476, RRID:CVCL_T841 | | | | | | |
| **Embryonic stem cells (control)** | | WA09, RRID:CVCL_9773 | | | | | | |
| **HEK 293T cells** | | ATCC Cat# CRL-11268, RRID:CVCL_1926 | | | | | | |
| **SH-SY5Y cells** | | ATCC, Cat# CRL-11266, RRID:CVCL_0019 | | | | | | |

Note: RRIDs, Research Resource Identifiers (RRIDs); M, Male; F, Female.

**Table S2 (related to Figures 1 and 5). Comparisons of diMNs and iPSC-MNs**

| | | diMNs | iPSC-MNs |
|---|---|---|---|
| | **Yield** | Medium | High |
| | **Purity** | Medium | High |
| | **Culture time to maturity** | At least 6 weeks | Within 4 weeks |
| **Phenotypes of DYT1 MNs** | Neurite outgrowth [1] | 63% | 75% |
| | Primary branches [1] | 58% | 69% |
| | mRNA export activity [1] | 55% | 63% |
| | Protein transport activity [1, 2] | 37% | 52% |
| | Nuclear morphology [3] | 56% | 39% |
| | Mislocalized LMNB1 [3] | 87% | 57% |

Note: MNs, motor neurons; diMNs, directly induced MNs; iPSC-MNs, induced pluripotent stem cell-derived MNs; [1] shown as percentage of healthy controls; [2] Average of protein import and protein export activities; [3] shown as percentage of DYT1 MNs with abnormalities.

**Table S3 (related to Figures 6-8). Sequences of probes, primers, and targets of shRNAs**

| Name | Sequence (5'-3') |
|---|---|
| Oligo-dA probe | AAAAAAAAAAAAAAAAAAAAAAAAA-Digoxigenin |
| Oligo-dT probe | TTTTTTTTTT TTTTTTTTTTTTTTT-Digoxigenin |
| qRTPCR-LMNB1-F | AGGAAAGCGGAAGAGGGTTG |
| qRTPCR-LMNB1-R | GCCTCCCATTGGTTGATCCT |
| TOR1A-shRNA-1 | AACGGTGTTCACCAAGTTAGAT |
| TOR1A-shRNA-2 | CAGCAAGATCATCGCAGAGAATA |
| LMNB1-shRNA-1 | AGCTTCTTGATGTAAAGTTA |
| LMNB1-shRNA-2 | CAGACTGTCATCAGAGATGAA |

**Table S4 (related to Figures 1-8). List of antibodies used in this study**

| Name | Research Resource Identifiers (RRIDs) | Host | Immunostaining | Western Blot |
|---|---|---|---|---|
| CHAT | Millipore Cat# AB144P, RRID:AB_2079751 | Gt | 1:200 | - |
| DIG | Sigma Cat# 11333089001, RRID:AB_514496 | Sh | 1:100 | - |
| GFP | Antibodies Incorporated Cat# GFP-1020, RRID:AB_10000240 | Ck | 1:1000 | - |
| GLE1 | Abcam Cat# ab96007,RRID:AB_10678755 | Rb | - | 1:2000 |
| HB9 | DSHB Cat# 81.5C10, RRID:AB_2145209 | Ms | 1:500 | - |
| LAP2A | Abcam Cat# ab5162, RRID:AB_304757 | Rb | - | 1:2000 |
| LAP2B | Thermo Fisher Scientific Cat# A304-840A, RRID:AB_2621035 | Rb | - | 1:2000 |
| LMNA/C | Abcam Cat# ab40567, RRID:AB_775967 | Ms | 1:300 | - |
| LMNB1 | Proteintech Group Cat# 12987-1-AP, RRID:AB_2136290 | Rb | 1:500 | 1:2000 |
| LMNB2 | Abcam Cat# ab97513, RRID:AB_10681013 | Rb | 1:250 | 1:1000 |
| MAB414 | BioLegend Cat# 902901, RRID:AB_2565026) | Ms | 1:1000 | - |
| MAP2 | Sigma-Aldrich Cat# M4403, RRID:AB_477193 | Ms | 1:500 | - |
| MAP2 | Abcam Cat# ab5392, RRID:AB_2138153 | Ck | 1:10000 | - |
| NESTIN | Millipore Cat# MAB353, RRID:AB_94911 | Ms | 1:400 | - |
| NUP98 | Santa Cruz Biotechnology Cat# sc-74578, RRID:AB_2157953 | Ms | 1:200 | 1:1000 |
| NUP358 (RANBP2) | * Millipore, ABN1385 | Rb | 1:100 | 1:500 |
| OCT3/4 | Santa Cruz Biotechnology Cat# sc-5279, RRID:AB_628051 | Ms | 1:50 | - |
| PAX6 | Sigma-Aldrich Cat# HPA030775, RRID:AB_10601243 | Rb | 1:500 | - |
| RANGAP1 | Cell Signaling Technology Cat# 36067, RRID:AB_2799093 | Rb | - | 1:1000 |
| SMI-32 | BioLegend Cat# 801701, RRID:AB_2564642 | Ms | 1:5000 | - |
| SOX2 | Santa Cruz Biotechnology Cat# sc-365823, RRID:AB_10842165 | Ms | 1:50 | - |
| SSEA-4 | DSHB Cat# MC-813-70 (SSEA-4), RRID:AB_528477 | Ms | 1:1000 | - |
| SYN1 | Cell Signaling Technology Cat# 5297, RRID:AB_2616578 | Rb | 1:200 | - |
| TOR1A | Cell Signaling Technology Cat# 2150, RRID:AB_2230690 | Ms | - | 1:1000 |
| TUBB3 | Covance Research Products Inc Cat# MMS-435P, RRID:AB_2313773 | Ms | 1:2000 | 1:10000 |
| TUBB3 | Covance Research Products Inc Cat# PRB-435P-100, RRID:AB_291637 | Rb | 1:2000 | - |
| β-Actin | Sigma-Aldrich Cat# A5441, RRID:AB_476744 | Ms | - | 1:60000 |

Note: Ck, chicken; Gt, goat; Ms, mouse; Rb, rabbit; Sh, sheep; - not determined in this study; * RRID is not available.
